# Platelet GARP-dependent activation of TGF-β1 limits inflammation and promotes cardiac repair after myocardial infarction

**DOI:** 10.64898/2026.07.01.735778

**Authors:** Cécile Dufeys, Julie Bodart, Audrey Ginion, Jérôme Ambroise, Noah Trusgnach, Estelle L. Ollivier, Caroline Bouzin, Davide Brusa, Camille Michiels, Yotis A. Senis, Zoltan Nagy, Alice Marino, Luc Bertrand, Christophe Beauloye, Sophie Lucas, Sandrine Horman

## Abstract

Platelets are increasingly recognized as active regulators of inflammation beyond their canonical hemostatic functions. Although platelets rapidly accumulate in the injured myocardium after myocardial infarction (MI), the mechanisms by which they coordinate the inflammatory response remain poorly understood. Glycoprotein A repetitions predominant (GARP) is a membrane receptor that presents latent transforming growth factor-β1 (TGF-β1) on activated platelets and supports its activation. Given the central role of TGF-β1 in inflammation and tissue repair, we hypothesized that platelet GARP-dependent activation of TGF-β1 regulates inflammatory resolution and repair after MI. Using mice with megakaryocyte-and platelet-specific *Garp* deletion, we demonstrate that loss of platelet GARP selectively impaired generation of bioactive TGF-β1 without altering platelet reactivity. Following permanent coronary artery ligation, platelet-specific *Garp* deficiency markedly increased mortality from ventricular rupture and exacerbated adverse left ventricular remodeling, independent of initial infarct size. Transcriptomic and histological analyses revealed heightened endothelial cell activation, increased leukocyte recruitment, delayed inflammatory resolution, and defective extracellular matrix deposition in the absence of platelet GARP. Mechanistically, platelet GARP-dependent TGF-β1 signaling restrained endothelial activation after MI. Together, these findings identify platelet GARP-mediated activation of TGF-β1 as a critical platelet-intrinsic counter-regulatory checkpoint that limits endothelial-driven inflammation and promotes infarct stabilization. Our study reveals an unexpected protective immunoregulatory function of platelets in cardiac repair after ischemic injury.

## Introduction

Myocardial infarction (MI) triggers a tightly orchestrated cascade of inflammatory and reparative responses that ultimately determines cardiac structural integrity and function [7, 22, 30]. Early sterile inflammation is initiated by the release of danger-associated molecular patterns from necrotic cardiomyocytes, leading to endothelial activation, cytokine production, and leukocyte recruitment into the infarcted myocardium [7, 22, 30]. Although this inflammatory phase is essential for the clearance of dead cells and tissue debris, its timely resolution is required for the transition to tissue repair, including cardiac fibroblast (CF) activation and extracellular matrix (ECM) deposition. Failure to properly coordinate this inflammatory-to-reparative switch promotes ECM disorganization, adverse left ventricular remodeling, and fatal cardiac rupture [7, 10, 22, 30].

Increasing evidence indicates that, beyond their canonical hemostatic functions, platelets actively regulate inflammatory responses through interactions with endothelial and immune cells [14, 39]. Platelets rapidly accumulate in the infarcted myocardium after injury [14, 20, 29, 38, 47, 49], yet their precise contribution to the orchestration of post-MI inflammation and repair remains incompletely understood. Platelets are the main source of circulating transforming growth factor-β1 (TGF-β1), which is stored in α-granules as a latent complex and released into the extracellular space upon platelet activation [23, 32, 36]. Latent TGF-β1 consists of mature TGF-β1 bound to the latency-associated peptide (LAP), which prevents binding of mature TGF-β1 to the cytokine receptor (TβR). To exert biological effects, latent TGF-β1 requires activation, a process consisting in a profound conformational change in LAP that releases mature TGF-β1 and allows its interaction with TβR [17, 18]. In the infarcted myocardium, TGF-β1 exerts time-dependent functions, limiting excessive early inflammation while promoting fibroblast activation and fibrotic repair [17, 18].

Glycoprotein A repetitions predominant (GARP) is a transmembrane protein that presents latent TGF-β1 at the cell surface and enables its activation. GARP-dependent TGF-β1 activation has been described in several cell types, including stimulated regulatory T lymphocytes (Tregs), megakaryocytes and platelets, stimulated B cells, endothelial cells, and hepatic stellate cells [31, 36, 44, 54]. However, the capacity of GARP to generate active TGF-β1 is cell- and context-dependent, and its functional relevance has been investigated predominantly in Tregs [4, 9, 11, 27, 28, 42]. In platelets, although GARP-dependent TGF-β1 activation has been reported not to influence thrombotic or hemostatic responses in mice [45], whether this pathway contributes to platelet-mediated regulation of sterile tissue injury remains unknown.

Given the central role of TGF-β1 in both inflammation and fibrotic repair, we hypothesized that that GARP-dependent activation of TGF-β1 by platelets regulates post-infarction healing [12, 18]. Using megakaryocyte- and platelet-specific *Garp* knockout mice, we demonstrate that platelet GARP is required for efficient TGF-β1 activation following MI. Loss of platelet GARP exacerbates endothelial activation and leukocyte recruitment, delays inflammatory resolution, impairs early ECM deposition, markedly increases susceptibility to ventricular rupture, and enhances adverse left ventricular remodeling. Together, these findings reveal a previously unrecognized platelet immunoregulatory pathway linking local TGF-β1 activation to cardiac repair after MI.

## Materials and methods

### Experimental animals

Male and female mice aged 10 to 12 weeks were housed under a 12-hour light/dark cycle with ad libitum access to standard chow and water. Animals were randomly assigned to experimental groups, and investigators performing outcome assessments were blinded to genotypes. Predefined exclusion criteria included perioperative death and absence of regional left ventricular wall motion abnormalities at day 1 after MI. Deaths occurring after postoperative recovery were included in survival analyses and assessed for evidence of ventricular rupture. Sample sizes were determined based on previous studies using this MI experimental mouse model [15].

### Murine model of MI

MI was induced by permanent ligation of the left anterior descending (LAD) coronary artery, as previously described [15]. Mice received buprenorphine (0.1 mg/kg) for analgesia 30 minutes before surgery and were anesthetized with ketamine (80 mg/kg) and xylazine (5 mg/kg). Mice were intubated, mechanically ventilated, and maintained at 37°C throughout the procedure. Following left thoracotomy at the fourth intercostal space, the LAD was ligated approximately 2 mm below the left auricle using an 8-0 polyamide suture. Sham-operated mice underwent identical procedures without LAD ligation.

### Echocardiography

Cardiac function and left ventricular remodeling were assessed at the indicated time points by transthoracic echocardiography in mice anesthetized with isoflurane, using a Vevo 3100 imaging system (VisualSonics) equipped with an MX550D transducer. Two-dimensional B-mode and M-mode images were acquired in long- and short-axis views. Infarct extent was assessed based on regional left ventricular wall motion abnormalities, as previously described [15].

### Platelets, plasma, and serum isolation and preparation

Mice were anesthetized with ketamine (100 mg/kg) and xylazine (10 mg/kg) and blood was collected by retro-orbital bleeding. For plasma and platelets, blood was collected into 1:6 volume acid-citrate-dextrose (Roth, C3821) containing 1 U/mL apyrase (Roth, A6132).

#### Platelets

Platelet-rich plasma (PRP) was obtained by sequential centrifugation (800 g for 5 seconds, followed by 100 g for 5 minutes). PRP was washed in acid-citrate-dextrose and centrifuged at 400 g for 5 minutes. Platelets were resuspended in modified Tyrode’s buffer (135 mM NaCl, 12 mM NaHCO_3_, 3 mM KCl, 0.3 mM Na_2_HPO_4_, 1 mM MgCl_2_, 5 mM D-glucose, 10 mM Hepes, 0.35% BSA, pH 7.4) at 2.5 × 10^5^ platelets/µL.

#### Platelet releasates

Platelets were stimulated with 0.3 µg/mL CRP (Cambcol) for 5 minutes at 37°C in the presence of 2 mM Ca²⁺. Releasates were collected after 15 seconds centrifugation at 13,000 g.

#### Plasma and serum

Plasma was centrifuged at 1,500 g for 15 minutes at 4°C. Serum was obtained from coagulated blood after 2 hours at room temperature and centrifugation at 2,000 g for 20 minutes. Samples were stored at -80°C.

### T- and B-cell preparation from splenocytes

Single-cell suspensions were prepared by mechanical dissociation of spleens, followed by filtration through 70 µm nylon filters, followed by red blood cell lysis (Thermo Fischer Scientific, 00433357). Cells were subsequently filtered through 40 µm nylon filter and resuspended in X-VIVO 15 medium (Lonza, 02-060Q) supplemented with Glutamax (Thermo Fischer Scientific, 35050061). T cells were stimulated for 24 hours using Dynabeads mouse T-activator CD3/CD28 (Thermo Fischer Scientific, 11456D). B cells were stimulated with a mixture containing 25 µg/mL F(ab’)2 fragment anti-mouse IgM (Jackson Immunoresearch, 115006075), 10 ng/mL mouse IL-21 (eBioscience, 14821162), 20 µg/mL anti-CD40, and 1 µM CpG oligonucleotides (InvivoGen, ODN2395). Before antibody staining, cells were incubated with anti-CD16/32 antibodies for 10 minutes at 4°C to block Fc receptors.

### Flow cytometry

#### Platelets

Washed platelets (250 x 10^5^/µL) were stimulated with 100 mU/mL thrombin (Roth, T6634) for 8 minutes at 37°C and incubated with the indicated antibodies or corresponding isotype controls for 20 minutes at room temperature with the following fluorescently conjugated antibodies: PE anti-GARP (1:12.5, Thermo Fischer Scientific, 12989182), APC anti-LAP (1:12.5, Biolegend, TW7-16B4), FITC anti-CD62p (1:12.5, BD Pharmingen, 553744), PE anti-activated αIIbβ3 (1:12.5, Emfret Analytics, MO23-2), and the corresponding isotype control.

#### Lymphocytes

Samples were incubated with the following fluorescently conjugated antibodies: PE-Cy7 anti-CD4 (1:200, Biolegend, 100422), PE anti-GARP (1:50, Thermo Fischer Scientific, 12989182), BV650 anti-B220 (1:100, Biolegend, 103241), FITC anti-CD8a (1:200, Biolegend, 100706), and the corresponding isotype controls in the presence of a viability dye (1:1,000, eBioscience, 65086514) for 30 minutes at 4°C. For intra-cellular staining, cells were fixed and permeabilized using Foxp3 Transcription Factor Staining Buffer (eBioscience, 00512343) during 20 hours at 4°C, then stained with APC anti-FOXP3 (1:100, eBioscience, 17577382) in the presence of anti-CD16/32 (1:100). Data were acquired using a FACSCanto II cytometer (platelets) or a LSR Fortessa cytometer (lymphocytes) (BD Biosciences) and analyzed using the FlowJo software (Tree Star).

### ELISA

Active and total TGF-β1 concentrations were measured in serum using the Mouse TGF-β1 DuoSet ELISA kit (R&D Systems, DY1679), according to the manufacturer’s instructions. Cardiac troponin I concentrations were measured in serum using a mouse cardiac troponin I ELISA kit (Life Diagnostics, CTNIUS), according to the manufacturer’s instructions.

### Histological analyses

#### Tissue processing

Hearts were fixed with 4% formaldehyde for 48 hours at room temperature, paraffin embedded (Tissue-Tek VIP6, Sakura), sectioned (5 µm thickness), and slices were deparaffinized and rehydrated.

#### Immunostaining

As previously described [37], antigen retrieval was performed in citrate buffer pH 5.7 (F4/80, PCNA, vimentin, αSMA, and CD31) or Tris-EDTA buffer pH 9 (CD41, pSmad, Ly6G, and CCR2). Tissue sections were blocked for 1 hour with 5% bovine serum albumin and incubated with primary antibodies overnight at 4°C (anti-pSmad1/2/3/5, 1:50, Abcam, Ab52903; anti-CD41, 1:250, Abcam, ab181582; anti-Ly6G, 1:2,000, BD pharmingen, 551459; anti-F4/80, 1:300, Cell Signaling, 70076; anti-CD31, Cell signaling, 77699; anti-CCR2, Thermo Fischer Scientific, MA5-41175) or 90 minutes at room temperature (anti-PCNA, 1:800, Cell Signaling, 13110; anti-vimentin, 1:10,000, Abcam, ab92547; anti-αSMA, 1:5000, Abcam, ab124964). Sections were subsequently incubated for 1 hour at room temperature with either an anti-rabbit (Envision+ System-HRP Labelled Polymer, Dako, K4003) or an anti-rat (Polymer HRP, OriGene, D35-18) secondary antibody. For immunohistochemistry, staining was developed for 5 minutes using 3,3′-diaminobenzidine (DAB) substrate (Dako, K3468). Sections were counterstained with hematoxylin for 5 minutes and mounted (Sakura, Tissue Tek Film). For immunofluorescence, sections were incubated for 10 minutes with tyramide reagents (1:200, Thermo Fischer Scientific). Iterative staining was performed when required. Nuclei were counterstained with Hoechst (1:1,000, Thermo Fischer Scientific, 62249), and slides were mounted (Dako, S3023).

#### Histochemical staining

For picrosirius red staining, sections were incubated with 2% phosphomolybdic acid for 2 minutes, washed with distilled water, and incubated in 0.1% picrosirius red solution for 2 hours at room temperature. Sections were then briefly rinsed in 0.01 M hydrochloric acid and distilled water. For Alcian blue staining, sections were incubated with 3% acetic acid for 3 minutes, stained with Alcian blue solution for 30 minutes at 37°C, briefly rinsed in 3% acetic acid, and counterstained with nuclear fast red for 5 minutes. For wheat germ agglutinin (WGA) staining, heart sections were washed with PBS and incubated for 2 hours with rhodamine-conjugated WGA (1:150, Vector laboratories, RL1022).

#### Slides digitizing and analysis

Slides were scanned using the Pannoramic SCANII slide scanner (3DHistech) (immunohistochemistry and histochemical staining) or the Axioscan.Z1 (Carl Zeiss) (immunofluorescence). Analyses were performed with the Visiopharm software.

### Western blot

Cells were lysed in buffer containing 50 mM Tris-HCl pH 7.5, 1 mM EDTA, 1 mM EGTA, 0.27 M sucrose, 1% Triton X-100, 20 mM glycerol-2-phosphate disodium, 50 mM NaF, 5 mM Na_4_P2O_7_.10H_2_O, 1 mM dithiothreitol, protease and phosphatase inhibitors (Thermo Fischer Scientific, 78446). Equal amounts of protein were separated by SDS-PAGE and transferred to PVDF membranes. Membranes were probed overnight at 4°C with antibodies targeting pSmad1/2/3/5 (1:2,000, Abcam, ab52903), GARP (1:1,000, R&D Systems, AF6229), or GAPDH (1:50,000, Cell Signaling, 5174). After 1 hour of incubation with corresponding secondary antibodies, bands were visualized using chemiluminescence (Merck, 11500694001) and quantified using ImageJ. Data are presented as normalized densitometric ratios relative to GAPDH.

### RNA isolation, RT-qPCR, and RNA sequencing

#### RNA isolation

Total RNA was isolated from cardiac tissue using the RNeasy Mini Kit (Qiagen, 74104) following phenol-chloroform lysis. Samples were treated with DNase (Qiagen, 79254) according to the manufacturer’s instructions.

#### RT-qPCR

Total RNA was reverse transcribed (iScript™ cDNA synthesis kit, Bio-Rad, 1708891) and qPCR was performed using SYBR Green (Eurogentec, RTSN1005NR) on a CFX Connect (Bio-Rad). *Rpl32* served as the housekeeping gene. Primer sequences are listed in Supplementary Table S1.

#### RNA sequencing

Library preparation and sequencing were performed by Biomarker Technologies BMK GENE (Münster, Germany). Briefly, the mRNA was isolated by Oligo(dT)-attached magnetic beads and randomly fragmented in fragmentation buffer. First-strand cDNA was synthesized with fragmented mRNA as template and random hexamers as primers, followed by second-strand synthesis with addition of PCR buffer, dNTPs, RNase H and DNA polymerase I. Purification of cDNA was processed with AMPure XP beads. Double-strand cDNA was subjected to end repair. Adenosine was added to the end and ligated to adapters. AMPure XP beads were applied here to select fragments within size range of 300-400 bp. cDNA library was obtained by certain rounds of PCR on cDNA fragments generated from step 4. In order to ensure the quality of library, Qubit 2.0 and Agilent 2100 were used to examine the concentration of cDNA and insert size. Q-PCR was processed to obtain a more accurate library concentration. Library with concentration larger than 2 nM is acceptable.

#### Bioinformatics

HISAT2 [24] was used to align the NGS sequences of each sample to the reference genome (Mus musculus.GRCm39.genome.fa) and to estimate the gene expression count matrix. Differential gene expression between pKO and control mice was performed using models based on the negative binomial distribution with the DESeq2 (v1.40.2), Bioconductor package [2]. Fold-changes associated with the genotype were adjusted for sex. Genes with an adjusted *P* value < 0.05 were considered differentially expressed. Gene set enrichment analysis was performed using fgsea (v1.24.0) with Hallmark, KEGG, and Gene Ontology Biological Process gene sets. Data have been deposited in the Gene Expression Omnibus (GEO) under accession number GSE324431.

### Human cardiac fibroblast culture

Human cardiac fibroblasts (HCF) (ScienCell Research Laboratories, 6300) were seeded at 20,000 cells/cm^2^ and cultured for 24 hours in HCF basal medium (Cell applications, 315500) supplemented with 10% serum (Cell applications, 316GS) and 0.1% antibiotics according to the manufacturer’s recommendations. Before treatment, cells were serum-starved for 4 hours in RPMI 1640 medium (Thermo Fischer Scientific, 11875093). HCF were then incubated with serum, plasma, or platelet releasate for 30 minutes and lysed for western blot analysis.

### Endothelial cell culture and platelet co-culture experiments

Human umbilical vein endothelial cells (HUVEC, Lonza, C2519A) were seeded at 30,000 cells/cm^2^ and cultured in EGM-2 (Lonza, CC-3156) supplemented with 2% serum (Lonza, CC-4176) and 0.1% antibiotics, according to the manufacturer’s recommendations. For recombinant TGF-β1 experiments, HUVEC were pretreated with 1 ng/mL recombinant TGF-β1 (Abcam, ab50036) for 6 hours before stimulation with 1.5 U/mL thrombin (Roth, T6634) for 2 hours. For co-incubation experiments, 112.5 x 10^5^ platelets/cm^2^ resuspended in RPMI 1640 were added for 5 hours before RNA extraction.

### Statistical analysis

Statistical analysis was performed using GraphPad Prism 10.1.0. All data herein are presented as the mean ± SEM. A P-value < 0.05 was considered statistically significant. Normality of continuous variables was assessed using the Shapiro-Wilk test, and homogeneity of variances was evaluated prior to parametric testing. For comparisons between two groups, two-tailed unpaired Student’s t-tests were used when data met normality assumptions; otherwise, the non-parametric Mann-Whitney test was applied. For multiple-group comparisons, one-way or two-way ANOVA followed by appropriate post hoc tests was performed as indicated. Survival analyses were conducted using the Kaplan-Meier method and compared by the log-rank (Mantel-Cox) test. Grubbs’s test was used to exclude statistical outliers

## Results

### Genetic deletion of *Garp* in platelets impairs active TGF-β1 release without affecting platelet reactivity

By crossing *Gp1ba* promotor-driven Cre recombinase transgenic mice (*Gp1ba*-*Cre^+/-^*) [34] with *Garp*^fl/fl^ mice [46], we generated megakaryocyte- and platelet-specific *Garp* KO mice (*Gp1ba*-*Cre^+/-^;Garp^fl/fl^*, pKO). *Gp1ba*-*Cre^-/-^;Garp^fl/fl^* littermates served as controls (WT) (Fig. 1a). pKO mice were viable, displayed normal hematologic parameters, and showed only a modest increase in mean platelet volume, consistent with previous reports of this Cre model [34] (Supplementary Table S2).

**Fig. 1.**
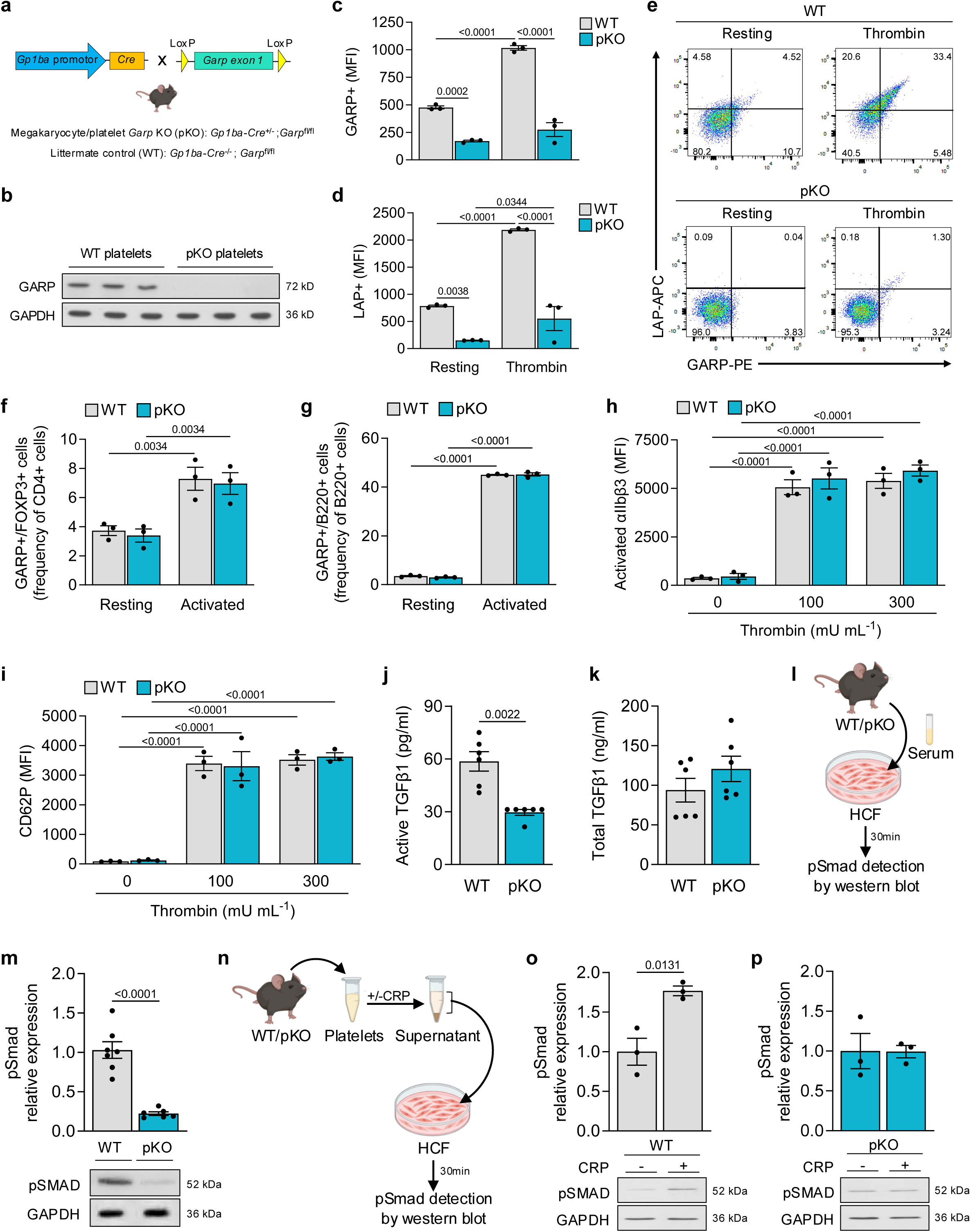
Genetic deletion of *Garp* in platelets impairs active TGF-β1 release without affecting platelet reactivity. **a** Schematic diagram depicting crossbreeding to generate a megakaryocyte/platelet-restricted GARP KO (pKO) mouse and its littermate control (WT). **b** Western blot analysis of GARP expression in lysates from WT and pKO platelets. GAPDH served as loading control (n=3 mice/group). **c-e** Flow cytometric analysis of surface GARP and LAP on WT and pKO platelets at rest and following stimulation with thrombin (100 mU/mL). Data are expressed as mean fluorescence intensity (MFI) (n=3 mice/group). **f,g** Flow cytometric analysis of GARP and FOXP3 expression on T lymphocytes (CD4+ cells) (**f**) and B lymphocytes (B220+ cells) (**g**) under resting and activated conditions. Data are expressed as frequency of CD4+ cells (**f**) or B220+ cells (**g**) (n=3 mice/group). **h,i** Surface expression of activated αIIbβ3 (**h**) and CD62P (**i**) on WT and pKO platelets at rest and following thrombin stimulation. Data are expressed as mean fluorescence intensity (MFI) (n=3 mice/group). **j,k** ELISA quantification of active (**j**) and total (**k**) TGF-β1 levels in the serum of WT and pKO mice (n=6 mice/group). **l** Experimental design for serum bioactivity assay shown in **m**. HCF were incubated for 30 minutes with serum of WT or pKO mice, and Smad phosphorylation in HCF was assessed by western blot. **m** Quantification of Smad phosphorylation in HCF following incubation with WT or pKO serum (n = 6-7 mice/group). **n** Experimental design for platelet releasate assay shown in **o,p**. HCF were incubated for 30 minutes with releasates from resting or CRP-activated WT or pKO platelets. **o,p** Smad phosphorylation in HCF following incubation with releasates from WT (**o**) or pKO (**p**) platelets (n = 3 mice/group). Data are presented as mean ± SEM. Statistical analysis was performed using unpaired two-tailed Student’s t-test (**m,o,p**), Mann-Whitney test (**j,k**), or two-way ANOVA followed by Fisher’s (**c,d,f,g**) or Tukey’s (**h,i**) multiple-comparisons test.

While expression of GARP was detectable in WT platelet lysates, it was markedly reduced in pKO platelets (Fig. 1b). Resting WT platelets expressed minimal levels of surface GARP and LAP which markedly increased upon activation with thrombin (Fig. 1c-e). In contrast, pKO platelets, whether resting or stimulated, exhibited very low levels of surface GARP expression (Fig. 1c-e). Expression of GARP on pKO Tregs and B lymphocytes, whether at rest or after T or B cell receptor stimulation, respectively, was similar to that of WT cells (Fig. 1f,g and Supplementary Fig. S1a,b). This confirmed the platelet-lineage specificity of the pKO. Integrin αIIbβ3 activation and surface CD62P exposure were similar between WT and pKO platelets following thrombin stimulation (Fig. 1h,i), indicating preserved platelet reactivity.

We next assessed whether platelet GARP deficiency alters TGF-β1 activation. Serum from pKO mice exhibited a ∼50% reduction in active TGF-β1, by comparison to WT, whereas concentrations of total TGF-β1 were unchanged (Fig. 1j,k). This indicates that release of active TGF-β1 in the serum, where platelets are fully activated and have discharged their granule contents, is drastically reduced in the absence of GARP on platelets. We confirmed this by measuring TGF-β1 activity in HCF incubated with serum from WT or pKO mice. Smad phosphorylation was measured as a readout of TGF-β1 activity (Fig. 1l). Serum from WT mice induced robust Smad phosphorylation in HCF, whereas serum from pKO did not (Fig. 1m). Consistently, releasates from activated WT platelets triggered Smad phosphorylation, while those from activated pKO platelets failed to do so (Fig. 1n-p).

Together, these findings establish that GARP is as a key determinant for production and release of active TGF-β1 by platelets but is not required for platelet activation.

### Platelet-specific deletion of GARP worsens post-MI outcome

To determine the functional relevance of platelet GARP during MI, WT and pKO mice were subjected to permanent ligation of the LAD coronary artery. Platelets rapidly accumulated within the infarcted myocardium in both genotypes. While early infiltration was comparable, platelet accumulation was significantly increased in pKO hearts by day 3 post-MI (Fig. 2a,b). The incidence of intramural platelet-rich thrombi did not differ between groups (Supplementary Fig. S2a,b). We measured levels of circulating active TGF-β1 by detecting Smad phosphorylation in HCF incubated with plasma collected from infarcted mice at day 1 post-MI. Plasma from infarcted WT mice induced robust Smad phosphorylation by comparison to sham controls, indicating increased systemic levels of active TGF-β1 on day 1 post-MI. This response was absent in plasma from pKO mice (Fig. 2c-e). These observations show that platelet GARP plays a critical role in the increase of circulating TGF-β1 activity following myocardial injury. Importantly, platelet GARP deficiency did not influence the extent of the initial ischemic insult. Plasma troponin I levels, and regional wall motion abnormalities were comparable between WT and pKO mice (Fig. 2f,g). Perioperative mortality was similar in both groups (WT: 10.4% vs pKO: 11.2%; n=125 per group). In contrast, mortality within the first 7 days after MI was markedly increased in pKO mice (Fig. 2h). Necropsy revealed a substantially higher incidence of fatal left ventricular wall rupture in pKO animals (38.5%) compared to WT controls (9.1%) (Fig. 2i,j). Among mice surviving the acute phase, pKO animals developed more pronounced adverse remodeling. At 14 days post-MI, pKO mice exhibited significantly increased left ventricular end-diastolic (LVED) and end-systolic (LVES) volumes (Fig. 2k,l) and diameters (Fig. 2m,n), with a trend toward a more reduced ejection fraction compared to WT mice (Table 1). Histological analysis confirmed a larger scar circumference in pKO hearts (Fig. 2o,p). In contrast, septal thickness, left ventricular mass, and cardiomyocyte cross-sectional area were comparable between genotypes (Table 1 and Supplementary Fig. S3a,b). Collectively, these observations reveal that platelet GARP is dispensable to control the magnitude of the initial ischemic injury, but it is essential to stabilize infarct and maintain structural integrity of the healing myocardium.

**Fig. 2.**
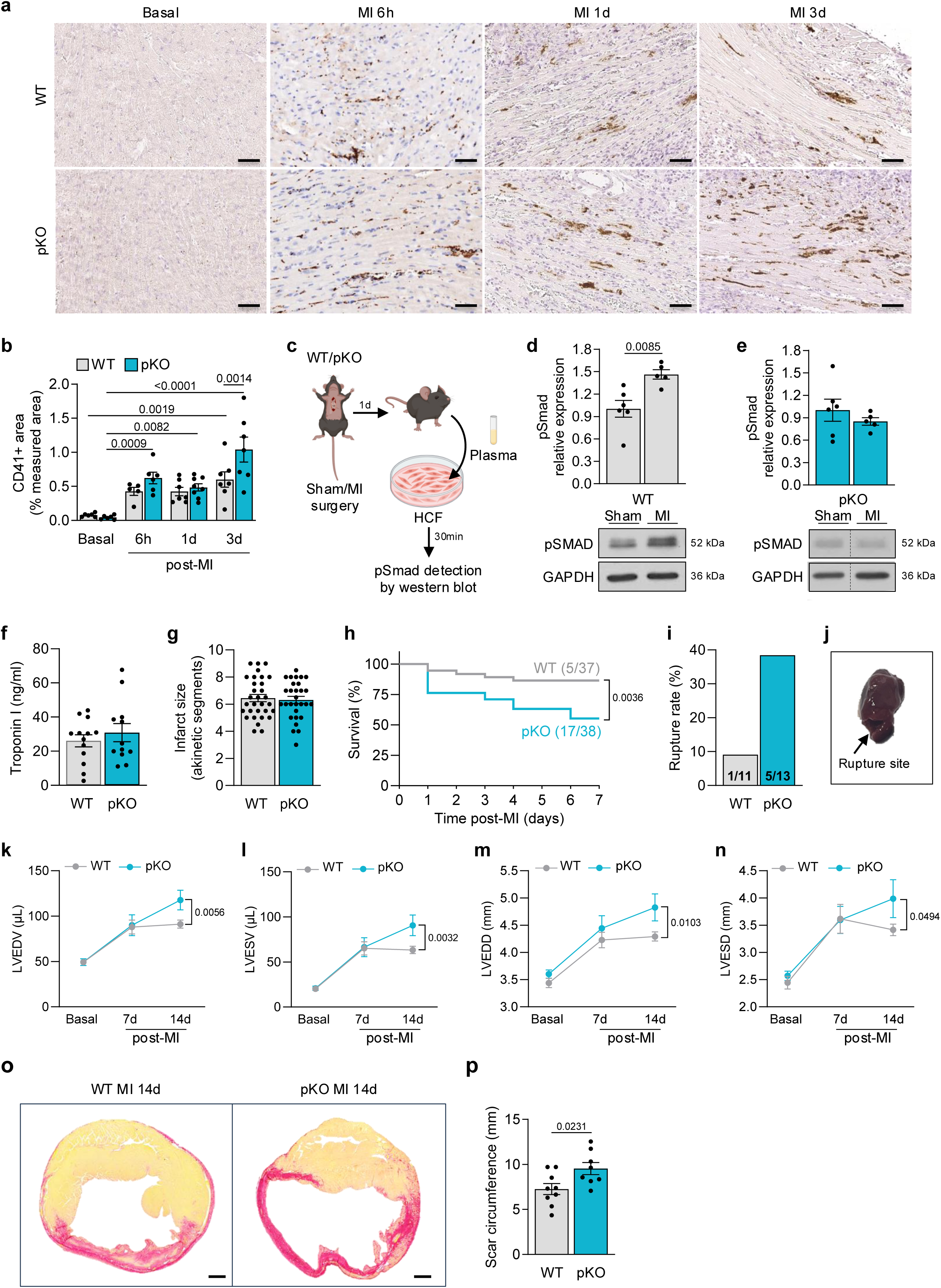
Platelet-specific deletion of GARP worsens post-MI outcome. **a** Representative images of platelets (CD41+) immunohistochemical staining in the myocardium of WT and pKO mice at baseline and at 6 hours, 1 day, and 3 days post-MI. Scale bar = 50 µm. **b** Quantification of CD41+ area shown in **a** (n=5-8 mice/group). **c** Experimental design for plasma bioactivity assay shown in **d,e**. HCF were incubated for 30 minutes with plasma collected from WT or pKO mice at 1 day after surgery, and Smad phosphorylation was assessed by western blot. **d,e** Smad phosphorylation in HCF following incubation with plasma isolated from WT (**d**) or pKO (**e**) mice at 1 day after surgery (n=5-6 mice/group). GAPDH served as loading control. **f** Plasma cardiac troponin I levels at 1 day post-MI (n=12-14 mice/group). **g** Echocardiographic assessment of infarct size at 3 days post-MI (n=29-31 mice/group). **h** Kaplan-Meier survival curves of WT and pKO mice during the first 7 days following MI (n=37-38 mice/group). **i** Incidence of left ventricle rupture in WT and pKO mice. **j** Representative image of left ventricle rupture in a pKO mouse. **k-n** Left ventricular end-diastolic volume (LVEDV, **k**), end-systolic volume (LVESV, **l**), end-diastolic diameter (LVEDD, **m**), and end-systolic diameter (LVESD, **n**) of WT and pKO mice measured by transthoracic echocardiography at baseline, 7 days, and 14 days post-MI (n=4-12 mice/group). **o** Representative images of picrosirius red staining of WT and pKO infarcted hearts at 14 days post-MI. Scale bar = 500 µm. **p** Quantification of scar circumference shown in **o** (n=8-9 mice/group). Data are presented as mean ± SEM. Statistical analyses were performed using unpaired two-tailed Student’s t-test (**d**-**g**,**p**), log-rank test (**h**), or two-way ANOVA followed by Tukey’s multiple-comparisons test (**b**,**k**-**n**).

**Table 1.** Echocardiographic parameters of WT and pKO hearts at baseline, 7 days, and 14 days post-MI. LVEDV, left ventricular end-diastolic volume; LVESV, left ventricular end-systolic volume; LVEDD, left ventricular end-diastolic diameter; LVESD, left ventricular end-systolic diameter; EF, ejection fraction; FS, fractional shortening; IVSd, end-diastolic interventricular septum thickness; IVSs, end-systolic interventricular septum thickness; PWd, diastolic posterior wall thickness; PWs, systolic posterior wall thickness; LV; left ventricular; TL, tibia length; CO, cardiac output; SV, stroke volume; HR, heart rate. Parameters are presented as mean±SEM. Statistical significance was determined by two-way ANOVA followed by Tukey’s multiple comparison test. P-value is represented as pKO compared to respective WT.

|  |  | WT | pKO | P value |
| --- | --- | --- | --- | --- |
| LVEDV (μl) | Basal | 49.86 ± 2.66 | 47.63 ± 3.19 | 0.9901 |
|  | MI 7 days | 87.96 ± 7.67 | 89.96 ± 11.49 | 0.8662 |
|  | MI 14 days | 91.19 ± 4.44 | 117.60 ± 10.79 | **0.0056\*** |
| LVESV (μl) | Basal | 20.16 ± 1.60 | 18.99 ± 2.11 | 0.9399 |
|  | MI 7 days | 65.05 ± 7.35 | 66.39 ± 10.52 | 0.9071 |
|  | MI 14 days | 63.32 ± 4.05 | 90.61 ± 11.42 | **0.0032\*** |
| LVEDD (mm) | Basal | 3.50 ± 0.08 | 3.50 ± 0.07 | 0.3695 |
|  | MI 7 days | 4.23 ± 0.14 | 4.44 ± 0.23 | 0.4146 |
|  | MI 14 days | 4.29 ± 0.08 | 4.82 ± 0.25 | **0.0103\*** |
| LVESD (mm) | Basal | 2.48 ± 0.10 | 2.42 ± 0.11 | 0.6231 |
|  | MI 7 days | 3.62 ± 0.27 | 3.60 ± 0.25 | 0.9543 |
|  | MI 14 days | 3.41 ± 0.11 | 3.99 ± 0.35 | **0.0494\*** |
| EF (%) | Basal | 60.50 ± 1.66 | 61.32 ± 2.52 | 0.7832 |
|  | MI 7 days | 27.34 ± 3.29 | 26.77 ± 3.98 | 0.9073 |
|  | MI 14 days | 30.73 ± 1.92 | 24.76 ± 3.02 | 0.1177 |
| FS (%) | Basal | 31.14 ± 1.93 | 30.68 ± 1.83 | 0.4600 |
|  | MI 7 days | 18.01 ± 3.45 | 19.28 ± 1.95 | 0.7816 |
|  | MI 14 days | 20.76 ± 1.16 | 18.24 ± 3.30 | 0.4659 |
| IVSd (mm) | Basal | 0.73 ± 1.16 | 0.78 ± 0.02 | 0.3667 |
|  | MI 7 days | 0.95 ± 0.09 | 0.81 ± 0.07 | 0.1042 |
|  | MI 14 days | 0.91 ± 0.03 | 0.93 ± 0.06 | 0.7988 |
| IVSs (mm) | Basal | 1.09 ± 0.04 | 1.12 ± 0.03 | 0.6862 |
|  | MI 7 days | 0.97 ± 0.11 | 1.16 ± 0.10 | 0.1367 |
|  | MI 14 days | 1.16 ± 0.04 | 1.13 ± 0.09 | 0.7662 |
| PWd (mm) | Basal | 0.64 ± 0.02 | 0.68 ± 0.02 | 0.1580 |
|  | MI 7 days | 0.71 ± 0.05 | 0.74 ± 0.05 | 0.6430 |
|  | MI 14 days | 0.78 ± 0.04 | 0.74 ± 0.03 | 0.4299 |
| PWs (mm) | Basal | 0.97 ± 0.03 | 1.07 ± 0.04 | 0.3109 |
|  | MI 7 days | 0.92 ± 0.06 | 0.92 ± 0.10 | 0.9709 |
|  | MI 14 days | 1.03 ± 0.04 | 0.97 ± 0.08 | 0.4586 |
| LV mass (mg) | Basal | 88.89 ± 2.30 | 98.06 ± 3.87 | 0.9709 |
|  | MI 7 days | 127.90 ± 8.69 | 113.00 ± 8.07 | 0.1009 |
|  | MI 14 days | 100.7 ± 3.32 | 106.6 ± 4.86 | 0.3904 |
| CO (mL/min) | Basal | 13.71 ± 0.61 | 13.42 ± 0.97 | 0.4958 |
|  | MI 7 days | 10.25 ± 1.16 | 11.84 ± 2.21 | 0.3735 |
|  | MI 14 days | 12.59 ± 0.59 | 12.28 ± 0.90 | 0.8242 |
| SV (μl) | Basal | 29.70 ± 1.35 | 28.64 ± 1.60 | 0.8405 |
|  | MI 7 days | 22.91 ± 2.20 | 23.57 ± 3.40 | 0.8536 |
|  | MI 14 days | 27.87 ± 1.75 | 27.02 ± 2.07 | 0.7572 |
| HR (bpm) | Basal | 473.40 ± 8.59 | 479.70 ± 10.11 | 0.8484 |
|  | MI 7 days | 472.00 ± 13.20 | 494.00 ± 22.70 | 0.2879 |
|  | MI 14 days | 456.00 ± 11.26 | 454.40 ± 6.49 | 0.9218 |

### Platelet-specific deletion of GARP amplifies early inflammatory programs after MI

To define early molecular consequences of platelet GARP deficiency following MI, we performed RNA sequencing on infarcted myocardium 1 day after permanent LAD ligation (Fig. 3a). Transcriptomic profiling revealed a marked modulation of inflammation-associated genes in pKO infarcted hearts compared to WT controls (Fig. 3b). Gene set enrichment analysis identified significant enrichment of pathways related to cytokine production and signaling, leukocyte migration, and cell-cell adhesion in pKO infarcts (Fig. 3c). Although individual genes within these pathways did not reach significance after stringent *P* value adjustment for transcriptomic multiple comparisons, several biologically relevant inflammatory candidates were identified (Fig. 3d). We therefore validated these candidates by RT-qPCR, which confirmed significant upregulation of key inflammatory mediators in pKO infarcted hearts, including *Il1b* and *Il6*, as well as neutrophil-attracting chemokines such as *Cxcl1* and *Cxcl5* (Fig. 3e). Genes associated with infiltrating myeloid populations, including *Il18rap* and *Mmp8*, were also increased (Fig. 3e), indicating enhanced inflammatory cell recruitment. Notably, transcripts indicative of endothelial cell activation, including *Selp*, *Sele*, *Serpin1*, and *Olr1* were upregulated in pKO hearts (Fig. 3e), supporting increased endothelial responsiveness in the absence of platelet GARP. In contrast, several genes remained unchanged (Supplementary Fig. S4a).

**Fig. 3.**
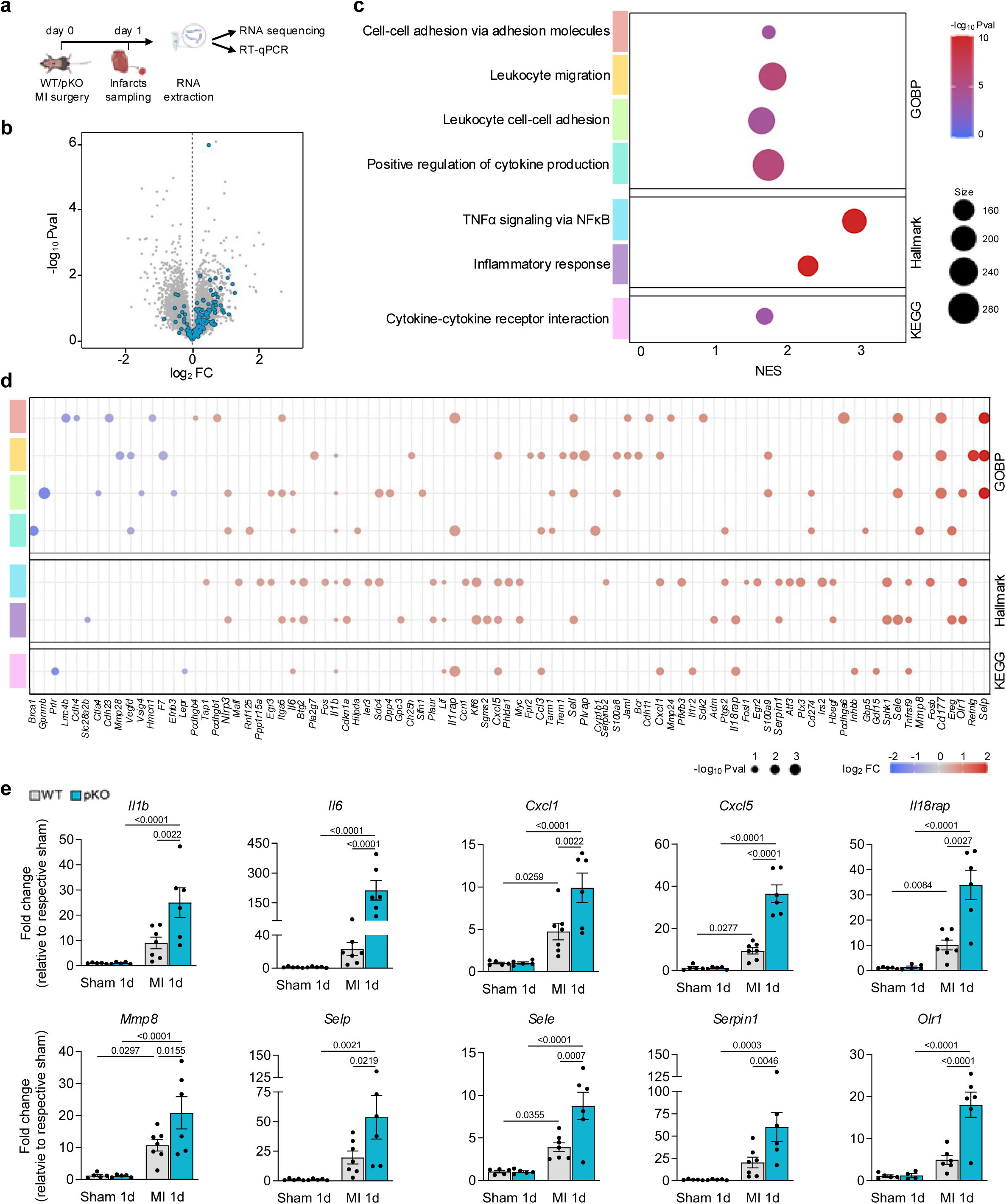
Platelet-specific deletion of GARP amplifies early inflammatory programs after MI. **a** Experimental design for Fig. 3. Infarcted myocardium was collected from WT and pKO mice at day 1 post-surgery. Total RNA was extracted and subjected to RNA sequencing analysis or RT-qPCR (n=6-8 mice/group). **b** Volcano plot showing differentially expressed genes in pKO versus WT infarcts. Blue dots highlight genes associated with inflammatory pathways. **c** Gene set enrichment analysis (GSEA) using Hallmark, Kyoto Encyclopedia of Gene and Genomes (KEGG), and Gene Ontology Biological Processes (GOBP) databases. Dot size represents the number of genes per pathway and color indicates -log10 *P* value. **d** Modulation of genes within enriched pathways. Dot size represents -log10 *P* value and color indicates log2 FC in pKO relative to WT infarcts. For the sake of clarity, only genes showing substantial modulation (|log_2 FC| > 1.5) are displayed. **e** RT-qPCR analysis of *Il1b*, *Il6*, *Cxcl1*, *Cxcl5*, *Il18rap*, *Mmp8*, *Selp*, *Sele*, *Serpin1*, and *Olr1* mRNA expression in sham and infarcted myocardium from WT and pKO mice at 1 day post-surgery (n=5-7 mice/group). P val, raw P value; FC, fold-change; NES, normalized enrichment score. Data are presented as mean ± SEM. Statistical significance was determined using two-way ANOVA followed by Fisher’s multiple-comparisons test (**e**).

Altogether, these insights indicate that platelet-specific deletion of GARP amplifies the early inflammatory programs following MI.

### Platelet-specific deletion of GARP enhances leukocyte recruitment and delays reparative remodeling after MI

Given the proinflammatory transcriptional profile observed in pKO infarcts, we next assessed immune cell infiltration following MI. Immunohistochemistry revealed minimal neutrophil (Ly6G+) and macrophage (F4/80+) presence at baseline, with no differences between genotypes (Fig. 4a-d). Following MI, neutrophil infiltration peaked at day 1, whereas macrophage accumulation was maximal at day 3. Both populations were significantly increased in pKO hearts compared to WT (Fig. 4a-d), indicating exaggerated inflammatory cell recruitment. Despite the established role of GARP TGF-β1 in Treg biology, infiltration by Tregs was comparable between WT and pKO mice at baseline and during the early post-MI period (Supplementary Fig. S5a-c), indicating that enhanced inflammation was not attributable to altered Treg accumulation.

**Fig. 4.**
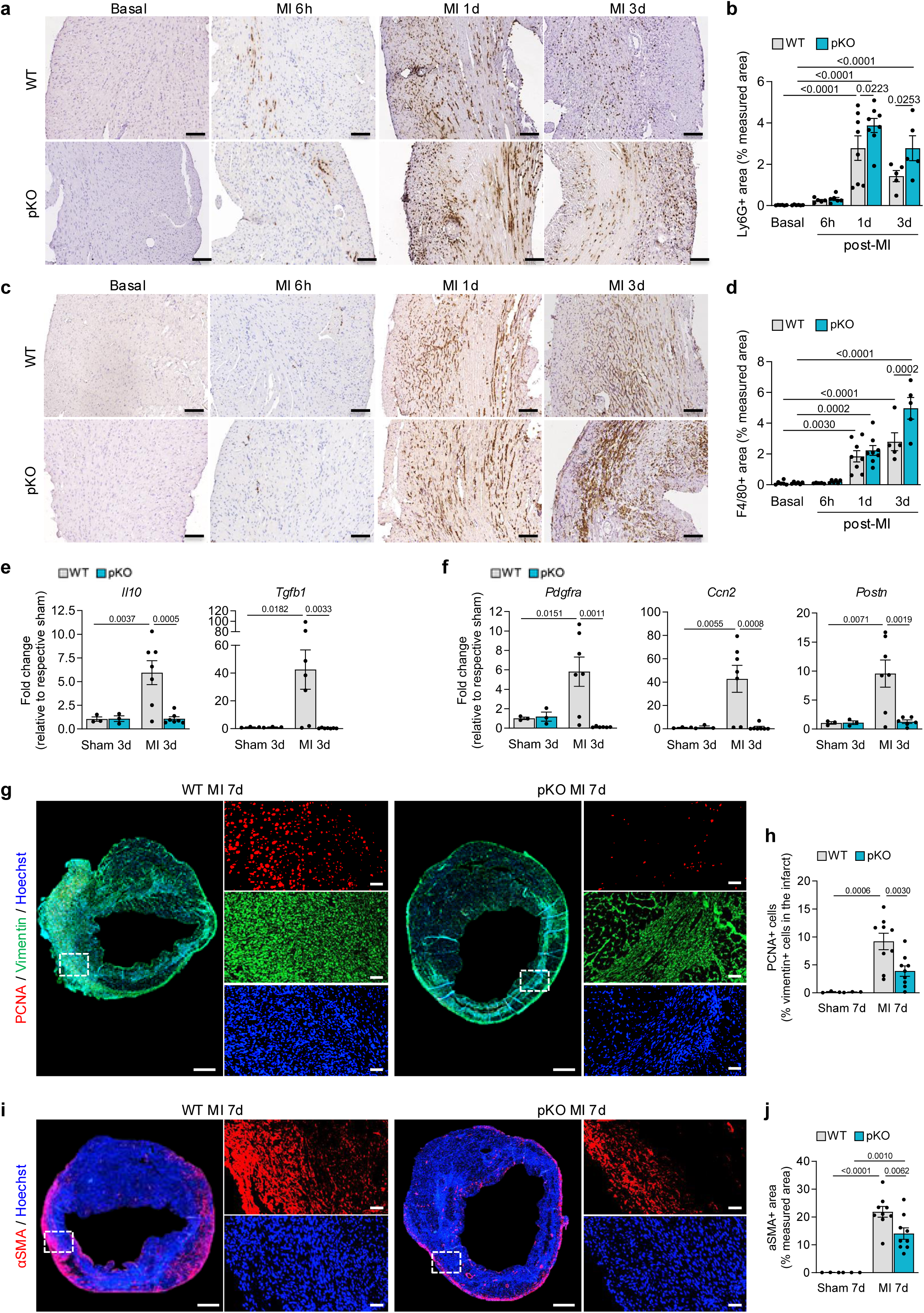
Platelet-specific deletion of GARP enhances leukocyte recruitment and delays reparative remodeling after MI. **a** Representative images of neutrophils (Ly6G+) immunohistochemical staining in the myocardium of WT and pKO mice at basal or at 6 hours, 1 day, and 3 days after MI. Scale bar = 100 µm. **b** Quantification of Ly6G immunostaining shown in a (n=5-8 mice/group). **c** Representative images of macrophages (F4/80+) immunohistochemical staining in the myocardium of WT and pKO mice at basal or at 6 hours, 1 day, and 3 days after MI. Scale bar = 100 µm. **d** Quantification of F4/80 immunostaining shown in **c** (n=5-8 mice/group). **e,f** RT-qPCR analysis of *Il10* and *Tfgb1* (**e**) or *Pdgfra*, *Ccn2*, and *Postn* (**f**) mRNA expression in the myocardium of WT and pKO mice at 3 days post-surgery (n=3-7 mice/group). **g** Representative images of proliferating CF co-immunohistofluorescent staining (PCNA: red, vimentin: green, nuclei: blue) in the myocardium of WT and pKO mice at 7 days post-surgery. Scale bar = 500 µm (left panel) or 50 µM (right panel). **h** Quantification of proliferating CF (PCNA/vimentin) co-immunohistofluorescent staining shown in **g** (n=3-9 mice/group). **i** Representative images of myofibroblasts immunohistofluorescent staining (αSMA: red, nuclei: blue) in the myocardium of WT and pKO mice at 7 days post-surgery. Scale bar = 500 µm (left panel) or 50 µM (right panel). **j** Quantification of myofibroblasts immunohistofluorescent staining (αSMA) shown in **i** (n=3-9 mice/group). Data are presented as mean ± SEM. Statistical significance was determined by two-way ANOVA followed by Fisher’s (**e**,**f**,**h**,**j**) or Tukey’s (**b**,**d**) multiple comparisons test.

We next evaluated the transition toward the reparative phase. Expression of the anti-inflammatory mediators *Il10* and *Tgfb1* was markedly reduced in pKO infarcts at day 3 post-MI (Fig. 4e), consistent with defective resolution of inflammation. In parallel, expressions of CF- and myofibroblast-associated genes, including *Pdgfra*, *Ccn2*, and *Postn*, were significantly decreased in pKO hearts (Fig. 4f). Histological analyses at day 7 post-MI confirmed reduced CF proliferation, as assessed by PCNA and vimentin co-staining, and diminished myofibroblast accumulation, as indicated by alpha smooth muscle actin (αSMA) staining (Fig. 4g-j). Overall, these findings confirm that platelet GARP deficiency amplifies early leukocyte recruitment and disrupts the inflammatory-to-reparative transition following MI.

### Platelet-specific deletion of GARP impairs ECM deposition after MI

Given the excessive inflammation and impaired transition to the reparative phase observed in pKO hearts, we next investigated the impact of platelet-GARP invalidation on ECM deposition. Histological analysis using alcian blue staining revealed a marked reduction in proteoglycan-and glycosaminoglycan-rich matrix in the infarcted myocardium of pKO mice compared to WT at day 3 post-MI (Fig. 5a,b). In addition, picrosirius red staining indicated a trend towards reduced collagen deposition in pKO infarcts (Fig. 5c,d). Consistently, induction of key ECM components at this time point, including *Fn1*, *Col1a1*, *Col3a1*, and *Sparc,* was robust in WT hearts after MI but significantly attenuated in pKO infarcts (Fig. 5e), indicating impaired early ECM synthesis. Moreover, expression of the neutrophil-derived collagenase *Mmp8* was markedly increased in pKO hearts (Fig. 5e), supporting enhanced ECM degradation mediated by inflammation-associated proteases. As angiogenesis accompanies ECM remodeling during infarct healing, we next assessed neovascularization by CD31 immunohistostaining. Vascular density increased over time in both genotypes, with no significant differences between WT and pKO mice (Supplementary Fig. S6a,b), indicating that impaired ECM deposition occurred independently of altered angiogenesis.

**Fig. 5.**
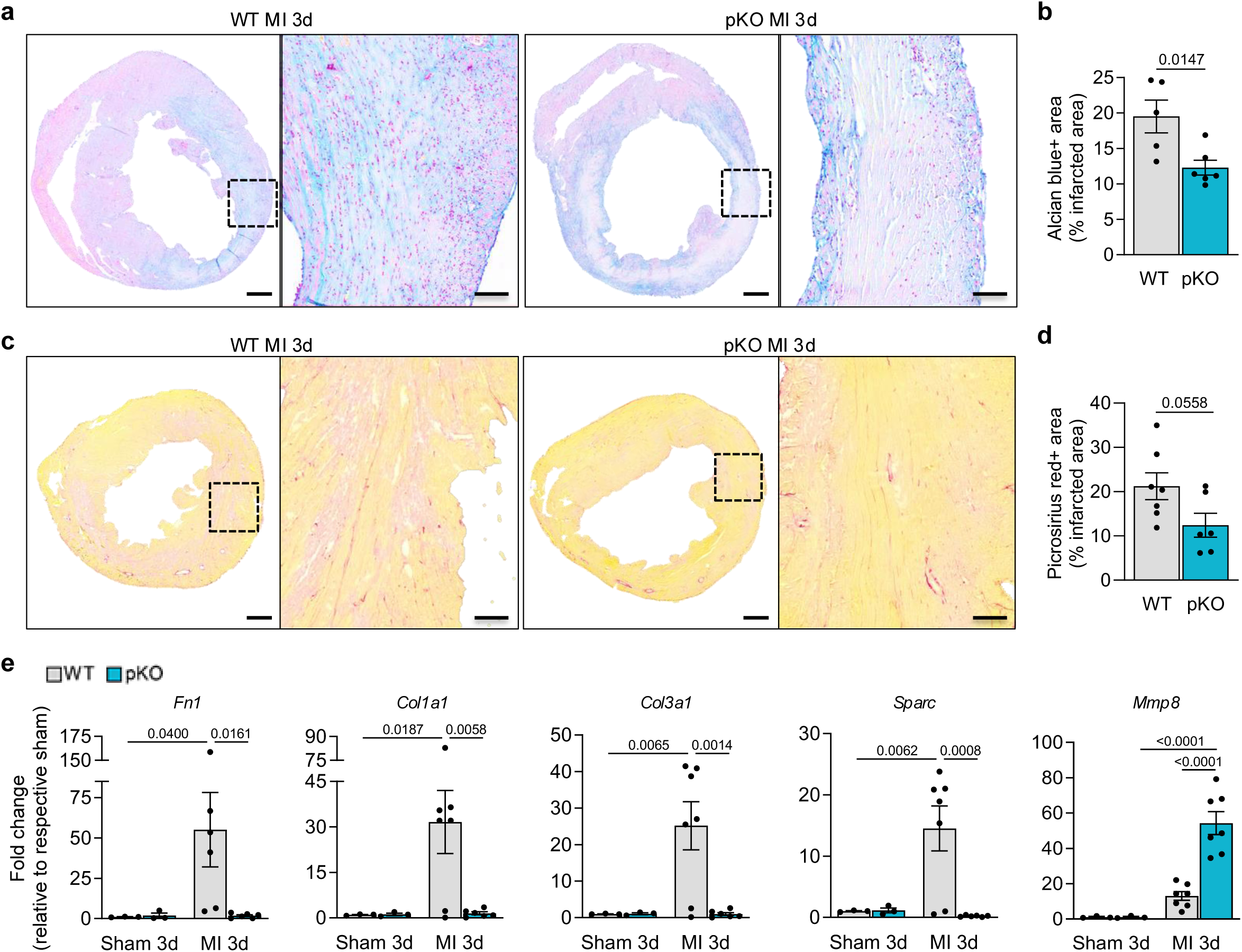
Platelet-specific deletion of GARP impairs ECM deposition after MI. **a** Representative images of alcian blue staining of WT and pKO infarcted hearts at 3 days post-MI. Scale bar = 500 µm (left panel) or 100 µm (right panel). **b** Quantification of alcian blue-positive area shown in **a** (n=6-7 mice/group). **c** Representative images of picrosirius red staining of WT and pKO infarcted hearts at 3 days post-MI. Scale bar = 500 µm (left panel) or 100 µm (right panel). **d** Quantification of picrosirius red-positive area shown in **c** (n=5-6 mice/group). **e** RT-qPCR analysis of *Fn1*, *Col1a1*, *Col3a1*, *Sparc*, and *Mmp8* mRNA expression in the myocardium of WT and pKO mice at 3 days post-surgery (n=3-7 mice/group). Data are presented as mean ± SEM. Statistical significance was determined using unpaired two-tailed Student’s t-test (**b**,**d**) or two-way ANOVA followed by Fisher’s multiple-comparisons test (**e**).

Taken together, these results provide a structural basis for the increased susceptibility to ventricular rupture and adverse remodeling observed in pKO mice.

### Platelet GARP-dependent TGF-β1 signaling restrains endothelial activation after MI

To define the cellular target of platelet GARP-dependent TGF-β1 signaling in vivo, we quantified nuclear Smad phosphorylation in neutrophils (Ly6G+), monocytes (CCR2+), and endothelial cells (CD31+) 6 hours after MI. Neutrophils exhibited low Smad phosphorylation, with no differences between WT and pKO (Fig. 6a,b). Monocytes displayed higher Smad activation, but again, levels were similar in WT and pKO hearts (Fig. 6c,d). In contrast, endothelial cells within the infarcted myocardium of pKO mice showed a marked reduction in nuclear Smad phosphorylation by comparison to WT mice (Fig. 6e,f), indicating a selective impairment of TGF-β1 signaling in the endothelium.

**Fig. 6.**
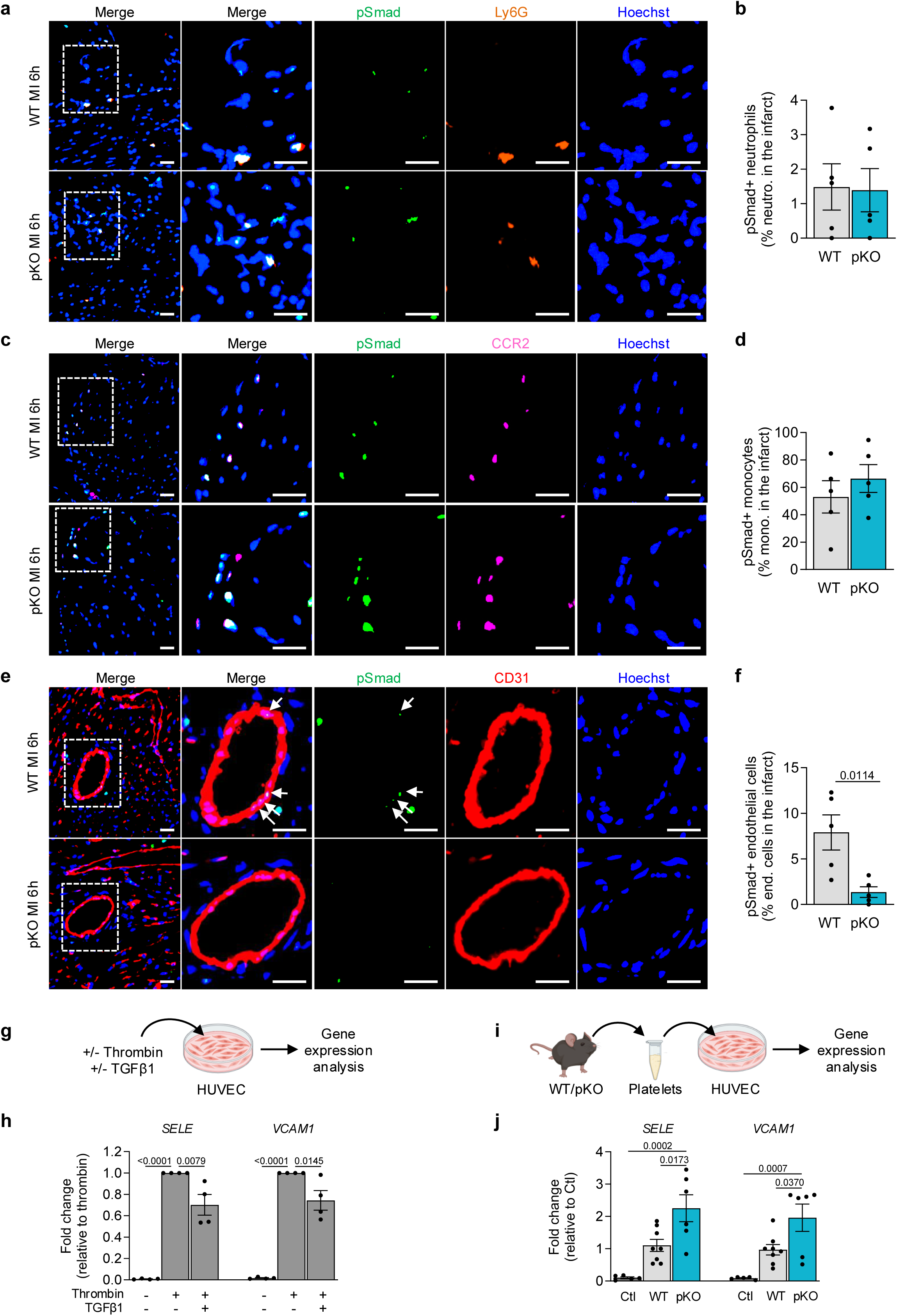
Platelet GARP-dependent TGF-β1 signaling restrains endothelial activation after MI. **a** Representative images of phosphorylated Smad (pSmad+, green) and neutrophils (Ly6G+, orange) co-immunohistofluorescent staining in WT and pKO infarcted myocardium at 6 hours post-MI. Scale bar = 25 µM. **b** Quantification of pSmad+ neutrophils shown in **a** (n=5 mice/group). **c** Representative images of phosphorylated Smad (pSmad+, green) and monocytes (CCR2+, pink) co-immunohistofluorescent staining in WT and pKO infarcted myocardium at 6 hours post-MI. Scale bar = 25 µM. **d** Quantification of pSmad+ monocytes shown in C (n=5 mice/group). **e** Representative images of phosphorylated Smad (pSmad+, green) and endothelial cells (CD31+, red) co-immunohistofluorescent staining in WT and pKO infarcted myocardium at 6 hours post-MI. Scale bar = 25 µM. **f** Quantification of pSmad+ endothelial cells shown in **e** (n=5 mice/group). **g** Experimental design for **h**. HUVEC were pretreated with recombinant active TGF-β1 (1 ng/mL) for 6 hours, followed by thrombin stimulation (1.5 U/mL) for 2 hours prior to gene expression analysis. **h** RT-qPCR analysis of *SELE* and *VCAM1* mRNA expression in HUVEC stimulated with thrombin in the presence or absence of recombinant active TGF-β1 (n=4 individual experiments). **i** Experimental design of **j**. HUVEC were incubated for 5 hours with platelets isolated from WT or pKO mice prior to gene expression analysis. **j** RT-qPCR analysis of *SELE* and *VCAM1* mRNA expression in HUVEC following stimulation with WT or pKO platelets (n=5-8 mice/group). Data are presented as mean ± SEM. Statistical significance was determined using unpaired two-tailed Student’s t-test (**b**,**d**,**f**) or one-way ANOVA followed by Dunnett’s (**h**) or Sidak’s (**j**) multiple-comparisons test.

We next examined whether TGF-β1 activity directly modulates endothelial cell activation, and whether reduced production of TGF-β1 by pKO platelets could perturb this activation. In cultured HUVEC, expression of adhesion molecules *SELE* and *VCAM1* was strongly induced by thrombin, and recombinant active TGF-β1 significantly attenuated this response (Fig. 6g,h). This demonstrates that TGF-β1 restrains endothelial cell activation. Interestingly, co-incubation of WT platelets with HUVEC only slightly induced the expression of *SELE* and *VCAM1* in the endothelial cells, whereas co-incubation with pKO platelets induced much higher *SELE* and *VCAM1* expression (Fig. 6i,j and Supplementary Fig. S7a,b). This indicates that reduced TGF-β1 production by pKO platelets fails to restrain endothelial cell activation. Collectively, our observations support a model in which platelet GARP is required for effective TGF-β1 signaling in endothelial cells after MI, and in which platelet-derived active TGF-β1 restrains endothelial activation, thereby limiting excessive leukocyte recruitment during the early inflammatory response (Fig. 7).

**Fig. 7.**
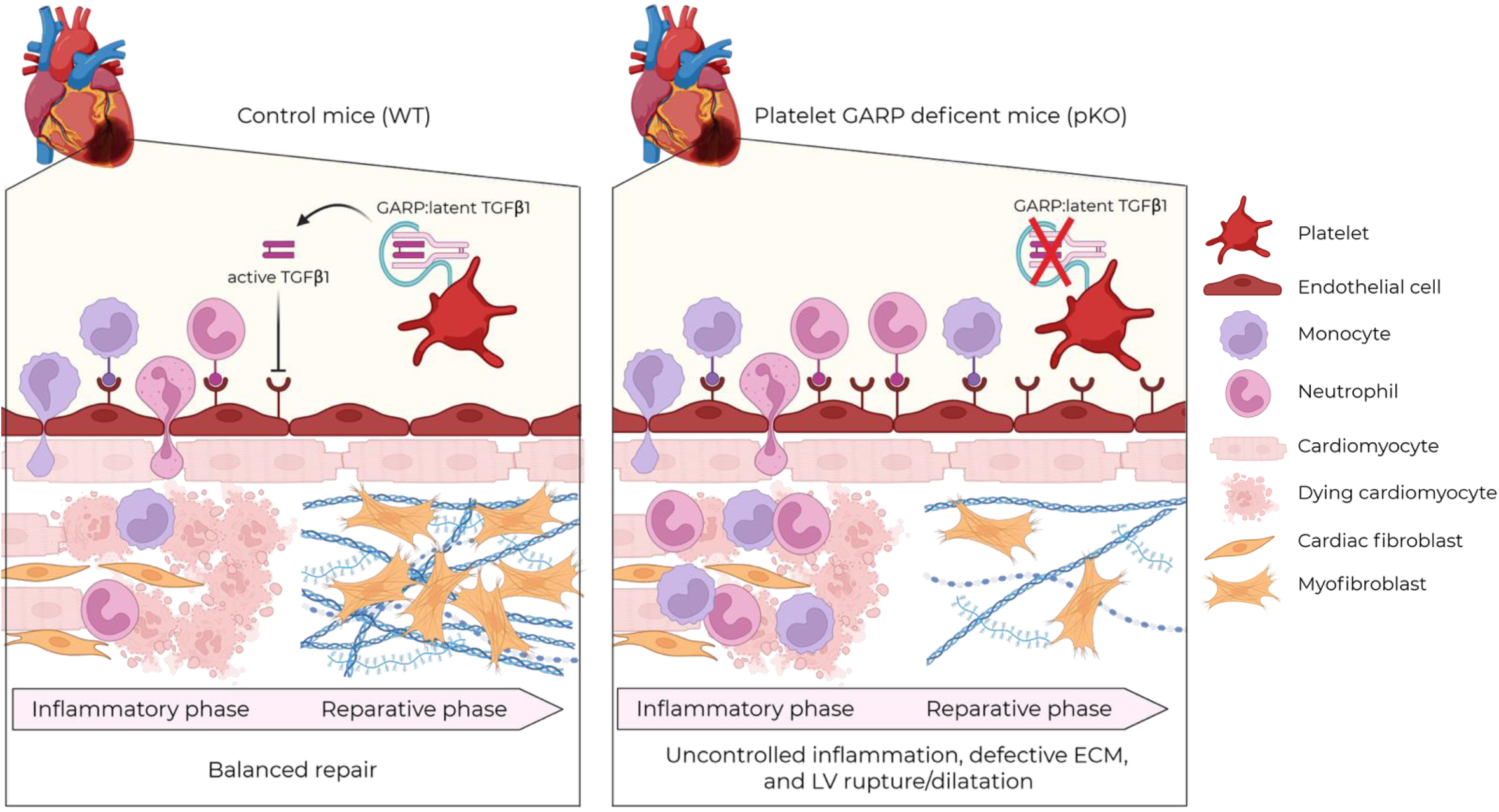
Activation of TGF-β1 by GARP-expressing platelets coordinates post-MI inflammation and repair. Schematic model illustrating the role of platelet GARP-dependent activation of latent TGF-β1 in post-MI healing. Following MI, platelet GARP promotes local TGF-β1 activation, thereby limiting endothelial adhesion molecule expression and ensuring controlled leukocyte recruitment. Loss of platelet GARP impairs platelet-dependent activation of TGF-β1, leading to excessive endothelial activation, enhanced inflammatory cell infiltration, defective fibroblast activation, and insufficient extracellular matrix deposition. This dysregulated reparative response increases susceptibility to left ventricular rupture and promotes adverse cardiac remodeling. Created with BioRender.com.

## Discussion

Despite major advances in reperfusion therapies that have reduced early mortality after MI, ventricular rupture and adverse remodeling remain major determinants of long-term outcome [5, 19]. Excessive or unresolved inflammation is a major driver of adverse cardiac remodeling and progression to heart failure after ischemic injury [19]. However, therapeutic strategies targeting broad inflammatory pathways, including IL1β or IL6 signaling, have yielded limited clinical benefits [25, 33]. These observations highlight the need to identify endogenous mechanisms that selectively restrain deleterious inflammation while preserving the reparative responses required for infarct stabilization. Accumulating evidence identifies platelets as active regulators of inflammation that act through direct cell-cell interactions and the secretion of a broad range of pro- and anti-inflammatory molecules [26, 39, 40, 43, 54]. Although platelets rapidly accumulate within the infarcted myocardium [14, 29, 38, 47, 49], the mechanisms by which these potentially protective functions contribute to inflammatory resolution and reparative remodeling remain poorly understood.

Using a megakaryocyte- and platelet-specific *Garp* knockout model, we identify platelet GARP as a critical regulator of post-MI healing through its control of TGF-β1 activation. Loss of platelet GARP selectively impaired generation of active TGF-β1 without altering platelet reactivity, indicating that this immunoregulatory function can be disrupted independently of conventional platelet activation responses. Platelet-specific GARP deletion exacerbated endothelial activation, increased expression of proinflammatory mediators and leukocyte recruitment, impaired fibroblast activation and early ECM deposition, and ultimately increased ventricular rupture and adverse remodeling despite a similar infarct size. Thus, platelet GARP-dependent TGF-β1 activation is not required for the initial extent of ischemic injury but is essential for stabilization of the healing infarct. These findings uncover a previously unrecognized platelet-dependent pathway controlling the transition from inflammation to reparative remodeling after MI.

A central mechanistic insight of our study is that endothelial cells emerge as a major early target of platelet GARP-dependent TGF-β1 signaling after MI. Endothelial Smad phosphorylation was markedly reduced in pKO infarcts, whereas signaling in neutrophils and monocytes was not detectably altered. Consistently, transcriptomic analyses revealed increased expression of endothelial activation markers and proinflammatory mediators involved in leukocyte recruitment in pKO infarcted hearts. Recent single-cell RNA-sequencing studies identified activated endothelial cells as important early sources of cytokines and chemokines, including *Il6* and *Cxcl1,* in the infarcted myocardium [1, 3], supporting the relevance of endothelial activation in the initiation and amplification of post-ischemic inflammation. *In vitro*, recombinant TGF-β1 attenuated thrombin-induced expression of adhesion molecules [6, 13, 21, 41, 48]. Platelets can themselves promote endothelial activation through the release of proinflammatory mediators [52]. Consistent with this property, co-culture with control platelets induced an increase in endothelial adhesion molecule expression. Importantly, this response was markedly exacerbated during co-culture with GARP-deficient platelets, indicating that platelet GARP-dependent TGF-β1 activation acts as an endogenous counter-regulatory mechanism limiting the proinflammatory effects of platelets on endothelial cells. This interpretation is also consistent with previous observations showing that monoclonal antibodies blocking GARP:TGF-β1 complexes enhance endothelial adhesion molecule expression in a murine tumor model [4]. Together, these observations identify the platelet GARP-TGF-β1 axis as an endogenous regulator of endothelial-driven inflammation after MI.

Platelet GARP deficiency also impaired the transition toward reparative remodeling. Reduced *Il10* and *Tgfb1* expression in pKO infarcts was associated with decreased fibroblast proliferation and impaired myofibroblast differentiation. These effects may result, at least in part, from the heightened inflammatory environment caused by reduced endothelial TGF-β1 signaling. However, a direct contribution of platelet-derived active TGF-β1 to fibroblast responses cannot be excluded and will require further investigation. Consistent with impaired fibroblast activation, early ECM deposition was reduced in pKO infarcts, providing a plausible mechanism for the increased susceptibility to ventricular rupture and dilation. Of note, neovascularization was not detectably altered in pKO infarcts, despite the established role of TGF-β1 in regulating angiogenesis [16]. Similarly, although platelet GARP-dependent TGF-β1 activation can modulate T cell function in the tumor microenvironment [36, 54], Treg accumulation was limited in the infarcted myocardium at the early time points examined after MI and was not affected by platelet GARP deletion. Thus, in this model, the early consequences of platelet GARP deficiency predominantly involve endothelial activation, innate immune cell recruitment, and infarct stabilization, without detectable effects on early Treg accumulation or neovascularization.

Several limitations should be acknowledged. First, *Gp1ba*-*Cre^-/-^;Garp^fl/fl^*littermates were used as controls, whereas *Gp1ba*-*Cre^+/-^;Garp^Wt/Wt^*mice were not included in the present study. Although platelet function and hemostatic responses have previously been reported to be largely preserved in *Gp1ba*-*Cre^+/-^*mice [35], a contribution of the Cre driver itself cannot be formally excluded. Second, our study relied on a permanent LAD ligation model, which provides a robust and reproducible framework to investigate inflammation, ECM deposition, infarct stabilization, and ventricular rupture, but does not fully reproduce the clinical scenario of ischemia-reperfusion injury. Whether platelet GARP-dependent TGF-β1 activation similarly regulates cardiac repair after reperfusion remains to be determined.

Our findings also raise important translational considerations. In acute MI, platelets are classically viewed primarily as drivers of thrombosis and inflammatory injury, providing the rationale for antiplatelet therapy. Our study indicates that platelets may also exert protective functions during infarct healing by limiting endothelial activation and inflammatory cell recruitment through GARP-dependent TGF-β1 activation. This functional duality should be considered when developing therapeutic strategies aimed at modulating platelet responses after ischemic injury. In particular, it will be important to determine whether antiplatelet agents commonly administered during acute MI affect platelet GARP expression or TGF-β1 activation, especially given increasing evidence that these therapies may exert context- and time-dependent immunomodulatory effects [8, 36, 50, 51, 53]. Conversely, antibodies blocking GARP:TGF-β1 complexes are currently being developed as cancer immunotherapies [4, 9, 11, 27, 28, 31]. Our findings raise the possibility that inhibition of this pathway may influence cardiac repair in the setting of ischemic injury. This potential effect warrants investigation in preclinical ischemia-reperfusion models and, ultimately, in translational and clinical studies.

In conclusion, our study reveals a protective immunoregulatory function of platelets during cardiac injury. Through GARP-dependent activation of TGF-β1, platelets restrain endothelial activation and excessive leukocyte recruitment, thereby promoting inflammatory resolution, infarct stabilization, and reparative remodeling after MI. Thus, beyond their established prothrombotic and proinflammatory activities, platelets can also engage endogenous anti-inflammatory pathways that preserve tissue integrity after ischemic injury. Collectively, these findings establish platelet GARP-dependent TGF-β1 activation as a key regulator of post-MI healing and highlight the need for future therapeutic strategies that selectively limit detrimental platelet activation while preserving beneficial platelet-mediated immune regulation.

## Declarations

### Ethics approval

Animal procedures were approved by the local authorities (Comité d’éthique facultaire pour l’expérimentation animale, 2021/UCL/MD/013) and conducted in accordance with the Guide for the Care and Use of Laboratory Animals (National Institutes of Health). The manuscript does not contain clinical studies or patient data.

## Acknowledgements

The authors would like to thank Emmanuel Vandenhooft for his dedicated animal care, and Aurélie Daumerie for her assistance with histological analyses.

## Author contributions

C.D., J.B., designed and conducted experiments, acquired and analyzed data, wrote, and edited the manuscript. A.G., C.M., N.T., E.O. conducted experiments. J.A. analyzed the transcriptomic data. Y.S., Z.N. provided *Gp1ba*-*Cre* mice. Ca.B., D.B. acquired and analyzed data. A.M., L.B. secured funding and edited the manuscript. Ch.B. designed research studies, secured funding, and edited the manuscript. S.L. provided *Garp^fl/fl^*mice and edited the manuscript. S.H. designed research studies, secured funding, wrote, and edited the manuscript. All authors read and approved the final manuscript prior to submission.

