## Supplementary Material for "Platelet GARP-dependent activation of TGF-β1 limits inflammation and promotes cardiac repair after myocardial infarction"

| Genes | Species | Forward | Reverse | T° |
|---|---|---|---|---|
| *Ccn2* | Mus musculus | CTGTGTACGGAGCGTGACCC | GACCCACCGAAGACACAGGG | 62.5°C |
| *Ccl3* | Mus musculus | GCCACATCGAGGGACTCTTCA | ATGGGGGTTGAGGAACGTGT | 64.4°C |
| *Cd177* | Mus musculus | GTGCAATGGAGCCGACAGCA | ACATCTCCTGGGGGAGGAACA | 60°C |
| *Col1a1* | Mus musculus | AGACGTGGAAACCCGAGGTA | TGGGTCCCTCGACTCCTA | 63.3°C |
| *Col3a1* | Mus musculus | GTCCACGAGGTGACAAAGGT | GATGCCCACTTGTTCCATCT | 60°C |
| *Cxcl1* | Mus musculus | CACTGCACCCAAACCGAAGTC | GGGAGCTTCAGGGTCAAGGC | 64.4°C |
| *Cxcl5* | Mus musculus | GCTGCCCCTTCCTCAGTCAT | ACTTCCACCGTAGGGCACTG | 64.4°C |
| *Fn1* | Mus musculus | GAGGGGACCCACAGTTCGTG | TCTTGCTCTTCCCGGCTTCG | 62.5°C |
| *Il1b* | Mus musculus | GCCACCTTTTGACAGTGATGAG | GACAGCCCAGGTCAAAGGTT | 64.4°C |
| *Il6* | Mus musculus | AAGAGTTGTGCAATGGCAATTCT | AAATTTTCAATAGGCAAATTTCCTGAT | 62.5°C |
| *Il10* | Mus musculus | GATGCCCCAGGCAGAGAA | CACCCAGGGAATTCAAATGC | 60°C |
| *Il1rap* | Mus musculus | CCGGCTCGAATCAAGTGCC | GATTCTCTGGGAGGCGGAAGT | 64.4°C |
| *Nlrp3* | Mus musculus | GCGCGTCTAGGTGAGAGTGT | CGCTCCTGCTTGCTTGGATG | 64.4°C |
| *Olr1* | Mus musculus | CTCCCCGTTCTGGATTGGAT | AGGTATGCACAGTTGCCTGA | 64.4°C |
| *Pdgfra* | Mus musculus | ACATCTGTGAGGCCACCGTC | AAGACGGCACAGGTCACCAC | 62.5°C |
| *Plvap* | Mus musculus | GTTGACTACGCGACGTGAGA | CTCGCTCAGGATGATAGCGG | 62.5°C |
| *Postn* | Mus musculus | ATGTTTATGGCACGCTGGGC | CAGGTTCTCCCAAGCCTCGT | 62.5°C |
| *Rpl32* | Mus musculus | GGCACCAGTCAGACCGATAT | CAGGATCTGGCCCTTGAAC | 60°c |
| *RPL32* | Homo sapiens | AGGCATTGACAACAGGGTTC | GTTGCACATCAGCAGCACTT | 62.5°C |
| *Sparc* | Mus musculus | GGAACAAATTCCGAGACGAA | TCTGAGCATCCTGCGTAATG | 60°C |
| *Serpin1* | Mus musculus | ACGTTGTGGAACTGCCCTAC | GCCAGGGTTGCACTAAACAT | 64.4°C |
| *Sele* | Mus musculus | AGAAAGCAAAGAAATTTGTTCCTGC | CCCACGATGCATTTGTGTTCCT | 62.5°C |
| *SELE* | Homo sapiens | GTTCAAGCCTGGCAGTTCCG | CACACAGTGCCAAACACGGG | 62.5°C |
| *Sell* | Mus musculus | GTGGACATGGGTGGGAACCAA | GCGTCATCGTTCCATTTCCCA | 64.4°C |
| *Selp* | Mus musculus | GGGGTTCGTGCTGAAGGGAA | CGAGCAGTTCACCTGGGCTT | 64.4°C |
| *Tgfb1* | Mus musculus | TTGCTTCAGCTCCACAGAGA | TGGTTGTAGAGGGCAAGGAC | 64.4°C |
| *VCAM1* | Homo sapiens | TTGACTTGCAGCACCACAGG | AGATGTGGTCCCCTCATTCGT | 62.5°C |

**Supplementary Table S1. Primers for RT-qPCR.** *Ccn2*, cellular communication network factor 2; *Ccl3*, C-C motif chemokine ligand 3; *Col1a1*, collagen type I; *Col1a3*, collagen type III; *Cxcl1*, C-X-C motif chemokine ligand 1; *Cxcl5*, C-X-C motif chemokine ligand 5; *Fn1*, fibronectin; *Il1b*, interleukin-1β; *Il6*, interleukin-6; *Il10*, interleukin-10; *Olr1*, oxidized low density lipoprotein receptor 1; *Pdgfra*, platelet-derived growth factor receptor alpha; *Plvap*, lasmalemma Vesicle-Associated Protein; *Postn*, Periostin; *Rpl32*, ribosomal protein L32; *Sparc*, Secreted Protein Acidic and Cysteine-Rich; *Sele*, E-selectin; *Sell*, L-selectin; *Selp*, P-selectin; *Tgfb1*, transforming growth factor-β1; *Vcam1*, vascular cell adhesion molecule-1.

| | WT | pKO | p-value |
|---|---|---|---|
| WBC (x$10^3$ cells/µl) | 4.60 ± 0.39 | 5.26 ± 0.36 | 0.2352 |
| Lymphocytes (%) | 84.03 ± 2.72 | 84.46 ± 1.53 | 0.8907 |
| Mid cells (%) | 9.64 ± 1.63 | 10.18 ± 1.27 | 0.7991 |
| Granulocytes (%) | 6.33 ± 1.13 | 5.37 ± 0.41 | 0.4293 |
| RBC (x$10^6$ cells/µl) | 7.56 ± 0.24 | 7.73 ± 0.17 | 0.5693 |
| Hemoglobin (g/L) | 11.00 ± 0.25 | 10.84 ± 0.18 | 0.6102 |
| Hematocrit (%) | 36.55 ± 0.76 | 35.22 ± 0.92 | 0.2819 |
| MCV (fL) | 53.00 ± 0.70 | 52.89 ± 0.57 | 0.9048 |
| Platelets (x$10^3$ cells/µl) | 731.30 ± 25.14 | 676.30 ± 18.91 | 0.0940 |
| MPV (fL) | 4.97 ± 0.06 | 5.55 ± 0.07 | **<0.0001** |

**Supplementary Table S2. Hematological parameters of WT and pKO mice at baseline.** Hematological parameters were analyzed with Cell-Dyn Emerald (Abbott Core Laboratory). WBC, white blood cells; RBC, red blood cells; MCV, mean cell volume; MPV, mean platelet volume. Parameters are expressed as mean ± SEM. Statistical significance was determined by unpaired t-test.

**Supplementary Fig. S1**

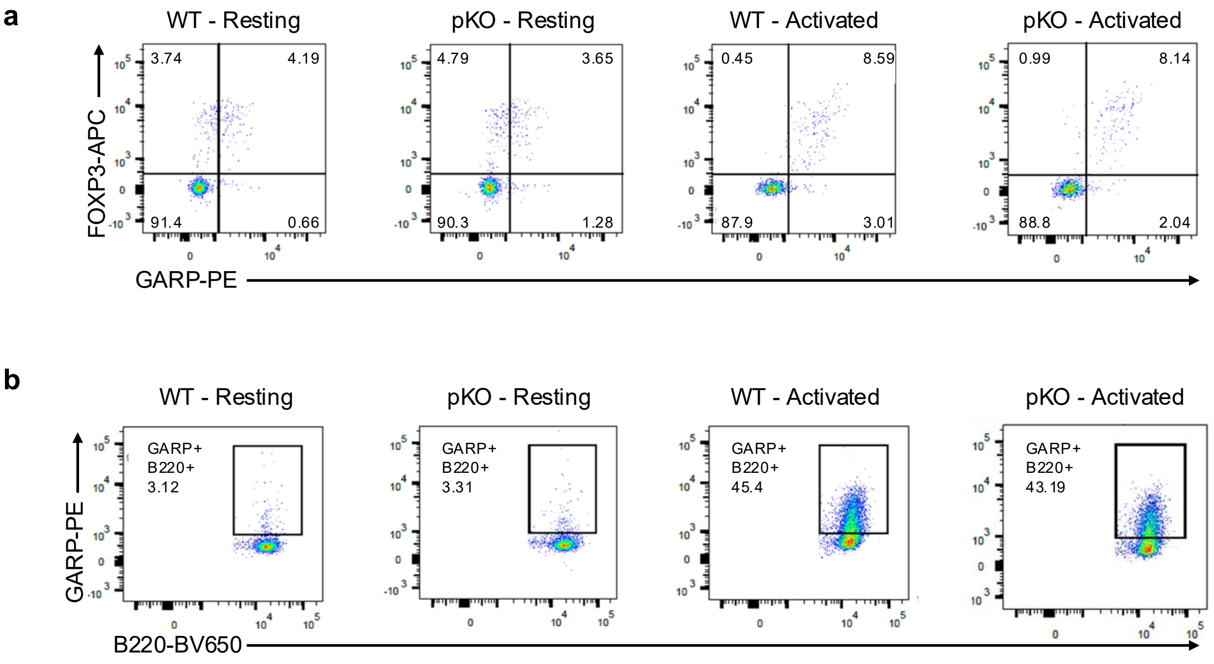


**Supplementary Fig. S1. Platelet-lineage specificity of genetic mouse model**
**a** Flow cytometry measurement of GARP and FOXP3 exposure on the surface of resting and activated CD4+ cells. **b** Flow cytometry measurement of GARP exposure on the surface of resting and activated B220+ cells.

**Supplementary Fig. S2**

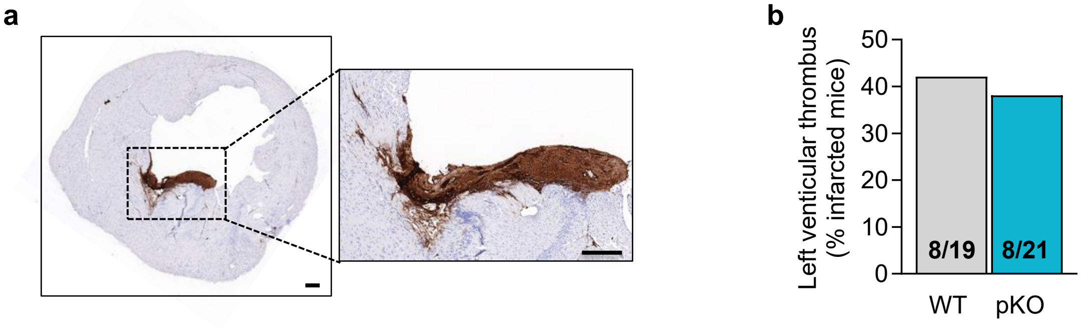


**Supplementary Fig. S2. Platelet-specific deletion of GARP does not impact left ventricular intramural thrombus formation after MI**

**a** Representative images of a left ventricular intramural thrombus in an infarcted heart, using platelets (CD41+) immunohistostaining. Scale bar = 250 µm. **b** Quantification of left ventricular intramural thrombus occurrence in WT and pKO infarcted mice (n=19-21 mice/group).

**Supplementary Fig. S3**

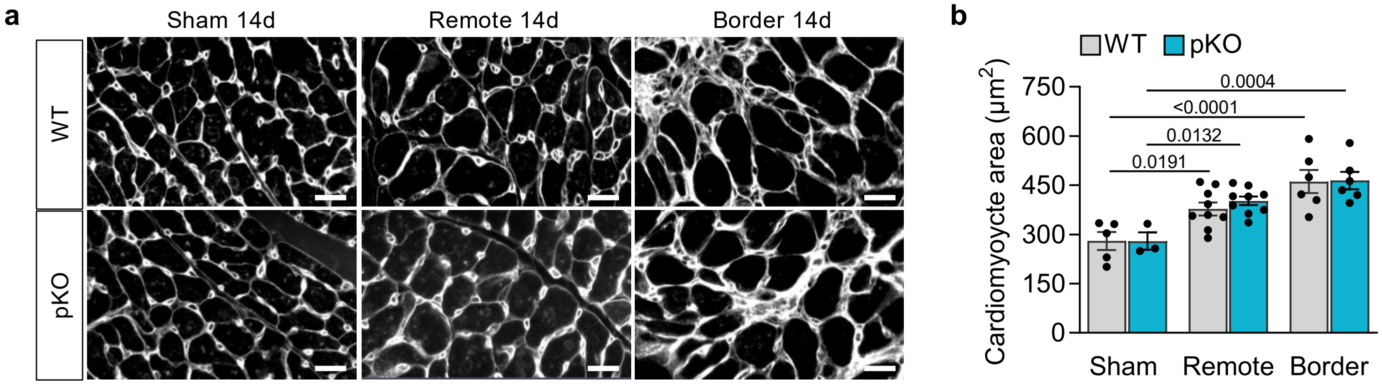


**Supplementary Fig. S3. Platelet-specific deletion of GARP does not impact cardiomyocyte hypertrophy after MI**

**a** Representative images of WGA staining in the sham, remote and border of WT and pKO myocardium, at 14 days post-surgery. Scale bar = 20 µm. **b** Quantification of cardiomyocyte area based on WGA staining shown in A (n=3-9 mice/group). Data are presented as mean ± SEM. Statistical significance was determined by two-way ANOVA followed by Tukey's multiple comparisons test .

**Supplementary Fig. S4**

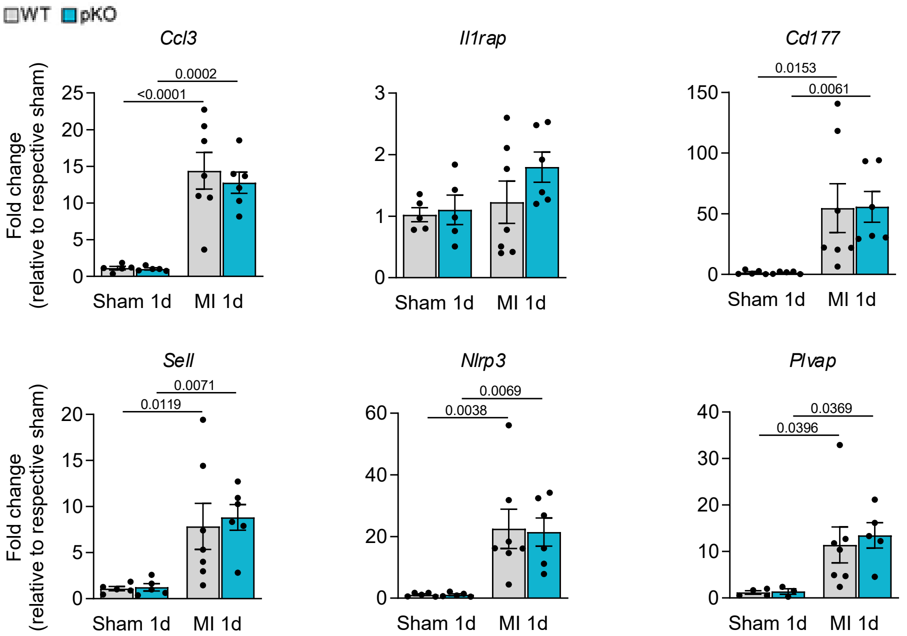


**Supplementary Fig. S4. , Non-differentially expressed genes identified in the differential expression analysis**

RT-qPCR measurement of *Ccl3*, *Il1rap*, *Cd177*, *Sell*, *Nlrp3*, and *Plvap* mRNA expression in sham and infarcted myocardium from WT and pKO mice at 1 day post-surgery (n=5-7 mice/group). Data are presented as mean ± SEM. Statistical significance was determined using two-way ANOVA followed by Fisher's multiple-comparisons test.

**Supplementary Fig. S5**

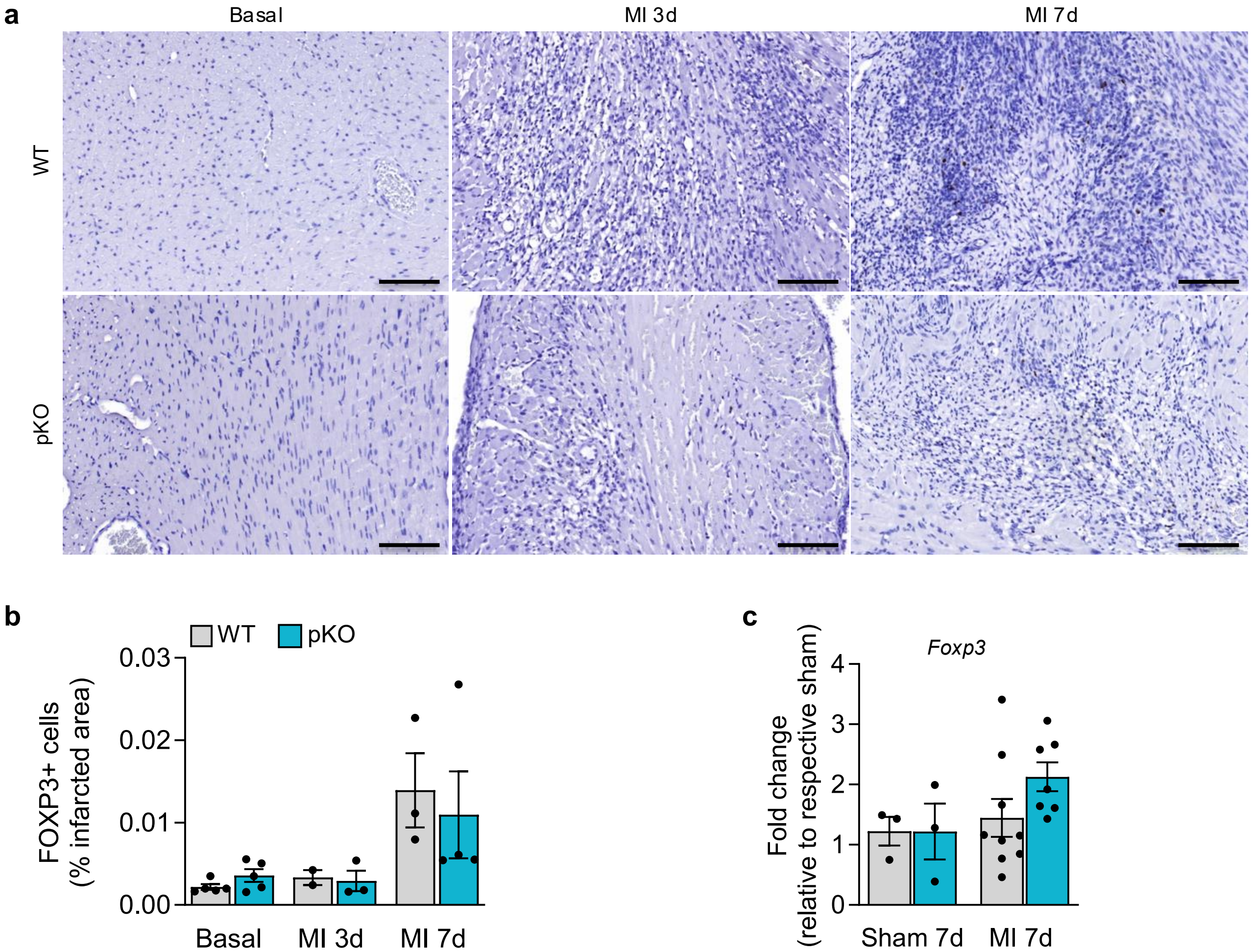


**Supplementary Fig. S5. Platelet-specific deletion of GARP does not affect Treg accumulation after MI**

**a** Representative images of Tregs (FOXP3+) immunohistochemical staining in sham or 3 days, and 7 days post-MI myocardium of WT and pKO mice. Scale bar = 100 µm. **b** Quantification of FOXP3 immunohistostaining shown in A (n=2-5 mice/group). **c** RT-qPCR analysis of *Foxp3* mRNA expression in the myocardium of WT and pKO mice at 7 days post-surgery (n=3-9 mice/group). Data are presented as mean ± SEM. Statistical significance was determined by two-way ANOVA followed by Tukey's (**b**) or Fisher's (**c**) multiple comparisons test .

**Supplementary Fig. S6**

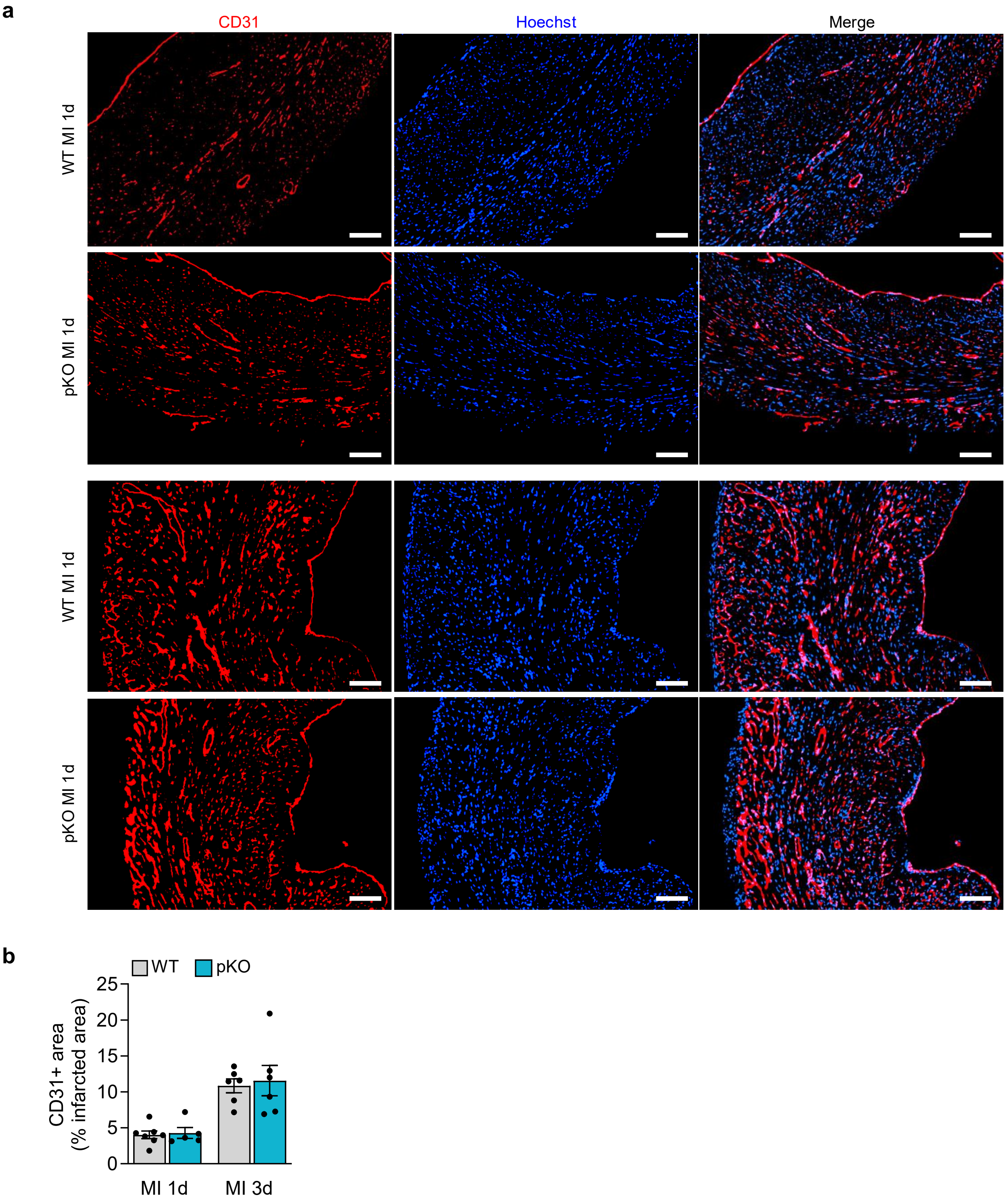


**Supplementary Fig. S6. Platelet-specific deletion of GARP does not affect angiogenesis after MI**
**a** Representative images of vessels (CD31+, red) immunohistofluorescent staining in WT and pKO infarcted myocardium at 1 day and 3 days post-MI. Scale bar = 100 μM. **b** Quantification of vessels (CD31+) immunohistofluorescent staining shown in **a** (n=6-7 mice/group). Data are presented as mean ± SEM. Statistical significance was determined by two-way ANOVA followed by Fisher's multiple comparisons test.

**Supplementary Fig. S7**

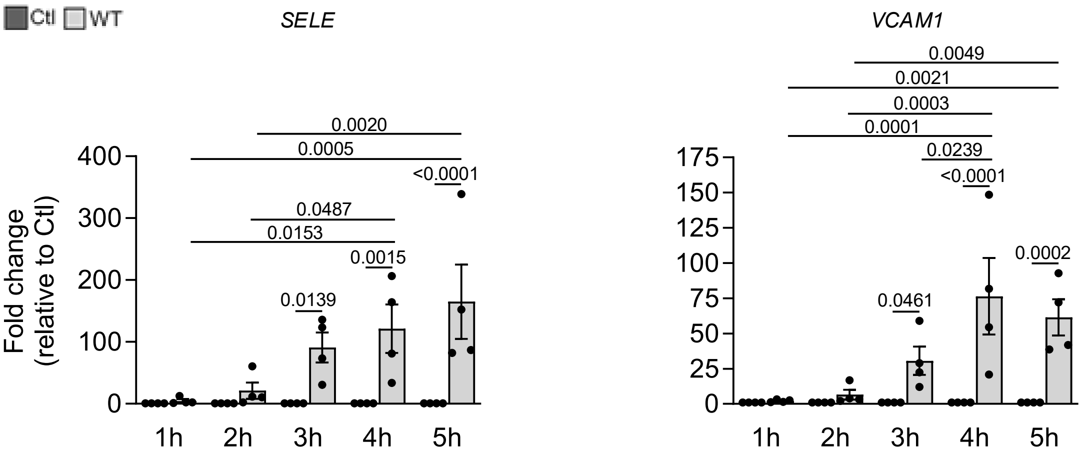


**Supplementary Fig. S7. Time-course of adhesion molecule expression in endothelial cells after co-incubation with WT platelets**

RT-qPCR analysis of *SELE* and *VCAM1* mRNA expression in HUVEC after increasing time of incubation with WT platelets. Data are presented as mean ± SEM. Statistical analyses were performed using two-way ANOVA followed by Tukey’s multiple-comparisons test.
